# Shared Song Detector Neurons in *Drosophila* Male and Female Brains Drive Sex-Specific Behaviors

**DOI:** 10.1101/366765

**Authors:** David Deutsch, Jan Clemens, Stephan Y. Thiberge, Georgia Guan, Mala Murthy

## Abstract

Males and females often produce distinct responses to the same sensory stimuli. How such differences arise – at the level of sensory processing or in the circuits that generate behavior – remains largely unresolved across sensory modalities. We address this issue in the acoustic communication system of *Drosophila*. During courtship, males generate time-varying songs, and each sex responds with specific behaviors. We characterize male and female behavioral tuning for all aspects of song, and show that feature tuning is similar between sexes, suggesting sex-shared song detectors drive divergent behaviors. We then identify higher-order neurons in the *Drosophila* brain, called pC2, that are tuned for multiple temporal aspects of one mode of the male’s song, and drive sex-specific behaviors. We thus uncover neurons that are specifically tuned to an acoustic communication signal and that reside at the sensory-motor interface, flexibly linking auditory perception with sex-specific behavioral responses.

## Introduction

Across animals, males and females produce distinct, dimorphic behaviors in response to common sensory stimuli (e.g., pheromones, visual cues, or acoustic signals), and these differences are critical for social and reproductive behaviors (Billeter and Levine, 2013; Dulac and Wagner, 2006; Kelley, 2003; Yamamoto et al., 2013; Yang and Shah, 2014). The molecular dissection of sexual dimorphisms in the nervous system of flies and mice in particular (Cachero et al., 2010; Dulac and Wagner, 2006; Rideout et al., 2010; Stowers and Logan, 2010; Yang and Shah, 2014; Yu et al., 2010) has identified neurons involved in either processing important social cues or driving social behaviors, but it remains open as to how sex-specific behaviors to common sensory signals emerge along sensorimotor pathways. It could be that males and females process sensory information differently, leading to different behavioral outcomes, or that males and females process sensory information identically, but drive different behaviors downstream of common detectors.

This issue has been most heavily investigated for pheromone processing. In *Drosophila*, the male pheromone cVA induces either aggression in males (Wang and Anderson, 2010) or receptivity in females (Billeter et al., 2009; Kurtovic et al., 2007). The pheromone is detected by shared circuits in males and females and the similarly processed sensory information (Datta et al., 2008) is then routed to sex-specific higher-order olfactory neurons (Kohl et al., 2013; Ruta et al., 2010) that likely exert different effects on behavior, although this hypothesis has not yet been tested. In the mouse, the male pheromone ESP1 triggers lordosis in females, but has no effect on male behavior. This pheromone activates V2Rp5 sensory neurons in both sexes but, analogous to cVA processing in flies, these neurons exhibit sex-specific projection patterns in the hypothalamus that drive sex-specific behavioral responses (Haga et al., 2010; Ishii et al., 2017). For pheromone processing then, the rule appears to be that early olfactory processing is largely shared between the sexes and then common percepts are routed to separate higher-order neurons or circuits for control of differential behaviors. But does this rule apply for other modalities, or for stimuli that can be defined by multiple temporal or spatial scales (e.g. visual objects or complex sounds)? For such stimuli, selectivity typically emerges in higher-order neurons (Dicarlo et al., 2012; Gentner, 2008; Tsao and Livingstone, 2008) and we do not yet know if such neurons are shared between males and females, and therefore if dimorphic responses emerge in downstream circuits.

Here, we investigate this issue in the auditory system in *Drosophila*. Similar to birds (Fortune et al., 2011; Konishi, 1985), frogs (Gerhardt and Huber, 2002), or other insects (Ronacher et al., 2014), acoustic communication in *Drosophila* involves different behaviors in males and females relative to the courtship song. During courtship, males chase females and produce a species-specific song that comprises two major modes – pulse song consists of trains of brief pulses and sine song consists of a sustained harmonic oscillation (Bennet-Clark and Ewing, 1967). In contrast with males, females are silent but arbitrate mating decisions (Bennet-Clark and Ewing, 1969). Males use visual feedback cues from the female (rapid changes in her walking speed and her distance relative to him) to determine which song mode (sine or pulse) to produce over time (Clemens et al., 2017; Coen et al., 2014; 2016) – this gives rise to the variable structure of song bouts (Fig. 1A). Receptive females slow in response to song (Aranha et al., 2017; Bussell et al., 2014; Clemens et al., 2015; Coen et al., 2014; Cook, 1973; Crossley et al., 1995; F. Von Schilcher, 1976; Tompkins et al., 1982), while playback of courtship song to males in the presence of other flies can induce them to increase their walking speed (Crossley et al., 1995; F. Von Schilcher, 1976; Vaughan et al., 2014), and to display courtship-like behaviors (Eberl et al., 1997; Li et al., 2018; Yoon et al., 2013; Zhou et al., 2015). These behavioral differences surrounding song production and perception between *Drosophila* males and females, combined with the wealth of genetic and neural circuit tools, make the *Drosophila* acoustic communication system an excellent one in which to investigate whether males and females share common sensory detection strategies for their courtship song, and how divergent behaviors arise.

**Figure 1.**
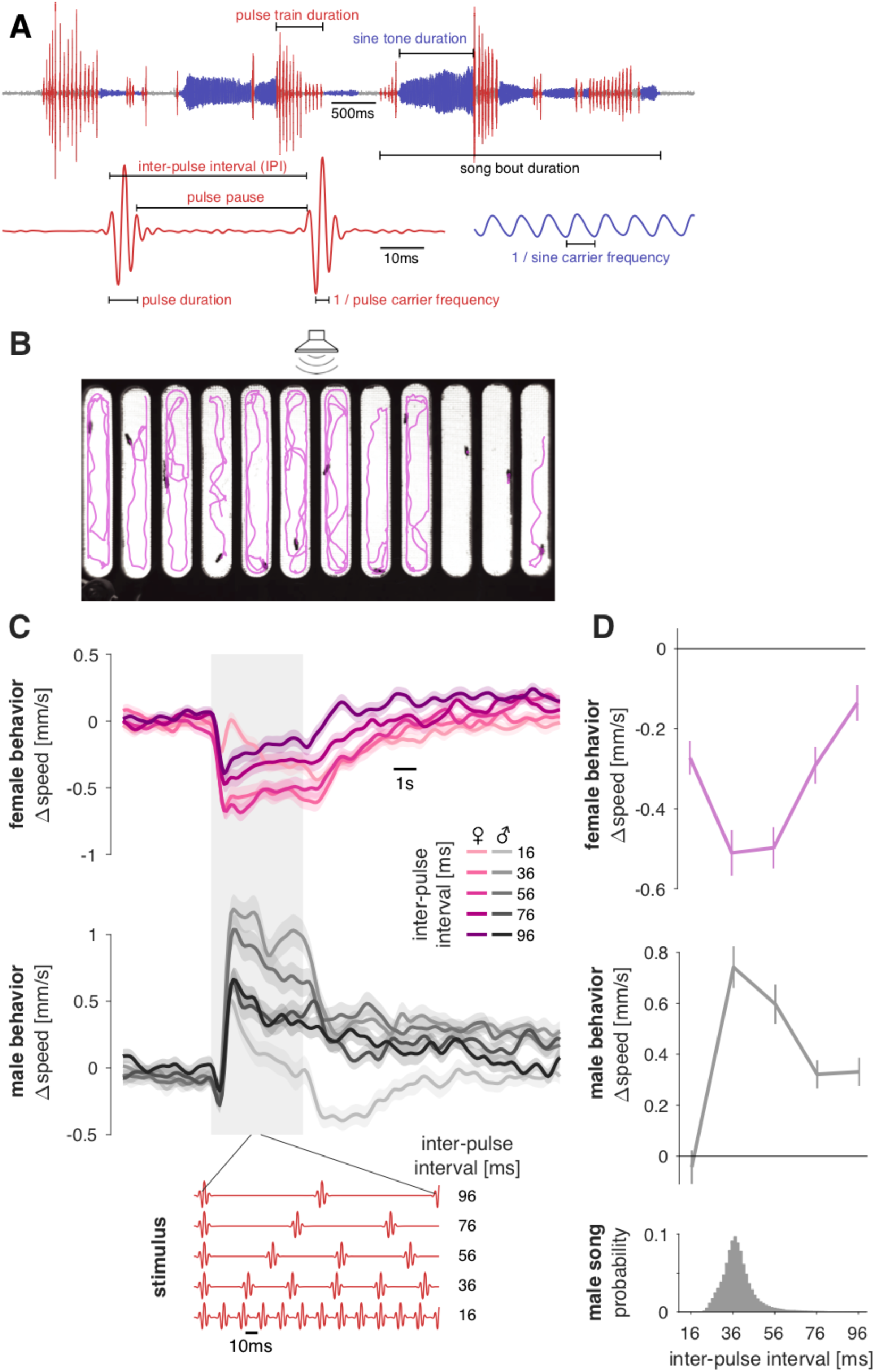
FLyTRAP assay for comparing locomotor tuning for courtship song stimuli in males and females. **A** *Drosophila melanogaster* produces song in bouts that can consist of two modes: Sine song corresponds to a weakly amplitude modulated oscillation with a species-specific carrier frequency (∼150Hz) and pulse song corresponds to trains of Gabor-like wavelets each with a carrier frequency between 220 and 450Hz and a duration between 6 and 12 ms. These pulses are produced at an inter-pulse interval (IPI) of 30-45 ms. **B** FLyTRAP consists of behavioral chamber that is placed in front of a speaker through which sound is presented. Fly movement is tracked using a camera. Shown is a single video frame of females in the assay with fly tracks for the preceding 20 seconds overlaid in magenta. See Movie S1. **C** Locomotor responses of females (magenta) and males (grey) for pulse trains with different IPIs (see legend). The gray shaded box indicates the duration of the sound stimulus. Red traces at the bottom of the plot show short snippets of the 5 IPI stimuli presented in this experiment. Baseline speed was subtracted before trial averaging. **D** Speed tuning curves for different IPIs in females (magenta) and males (grey) are obtained by averaging the speed traces in the six seconds following stimulus onset. The histograms at bottom shows the IPI distribution found in male song (data from 47 males of NM91 wild type strain totaling 82643 pulses). Lines and shaded areas or error bars in C and D correspond to the mean ± s.e.m. across 112 male and 112 female flies. All Δspeed values from the wild type strain NM91. See also Figure S1 and Movie S1.

Each major mode of *Drosophila* courtship song, sine or pulse, contains patterns on multiple temporal scales (Arthur et al., 2013; Bennet-Clark and Ewing, 1967) (Fig. 1A) – neurons that represent either the pulse or sine mode should in theory bind all of the temporal features of each mode, similar to object detectors in other systems (Bizley and Cohen, 2013; Dicarlo et al., 2012; Gentner and Margoliash, 2003; Griffiths and Warren, 2004), and their tuning should match behavioral tuning. Historically, behaviorally relevant song features have been defined based on the parameters of the species’ own song (Bennet-Clark and Ewing, 1969). However, there is now ample evidence that the preferred song can diverge from the conspecific song (Amézquita et al., 2011; Blankers et al., 2015; Ryan et al., 2001) – for instance if females prefer exaggerated song features (Rosenthal and Ryan, 2011; Ryan and Cummings, 2013) or respond to signal parameters not normally produced by their male conspecifics (Hennig et al., 2016). It is therefore important to define song modes by the acoustic tuning of specific behavioral outputs. This has been done for other insects (e.g. (Clemens and Hennig, 2013; D. von Helversen and O. von Helversen, 1997)) but never for flies in a systematic way that also permits a direct comparison between sexes.

To that end, we developed a behavioral assay for assessing dynamic changes in walking speed in response to sound playback in both sexes, and we then measured locomotor tuning for all features of either pulse or sine song. We found that males and females have similar tuning but different behavioral responses and that they are tuned for every major feature of the song. We then identified a small set of sexually dimorphic neurons, termed pC2 (Kimura et al., 2015; Rideout et al., 2010; Zhou et al., 2014), that serve as shared pulse song detectors in both sexes: the tuning of pC2 neurons is matched to behavioral tuning for pulse song – but not for sine song – across a wide range of temporal scales. We find that optogenetic activation of pC2 is sufficient to drive sex-specific behaviors – changes in locomotion with sex-specific dynamics as well as singing in males – and that silencing pC2 neurons biases males to production of sine song. pC2 is therefore both sensory and motor with regard to pulse song – it is important both for pulse song processing and pulse song generation. Finally, we establish the importance of pC2 neurons by showing that early social experience changes both the tuning of these neurons and the tuning of the behavior. Our results indicate that the fly brain contains common pulse song detectors in males and females which control sex-specific behavioral responses to song via downstream circuits.

## Results

### Comprehensive characterization of behavioral tuning for courtship song features

We designed a single-fly playback assay in which individual males or females receive acoustic stimuli in the absence of any confounding social interactions, and we implemented an automated tracker to analyze changes in locomotion relative to acoustic playback (Fig. 1B and Movie S1). The assay (which we refer to as FLyTRAP (**<u>F</u>**ly **<u>L</u>**ocomotor **<u>TR</u>**acking and **<u>A</u>**coustic **<u>P</u>**layback)) monitors dynamic changes in walking speed, which provides a readout that can be directly compared between both males and females, as opposed to slower readouts of sex-specific behaviors such as the female time to copulation (Bennet-Clark and Ewing, 1969; Zhou et al., 2014) or male-male chaining (Yoon et al., 2013; Zhou et al., 2015). Because of the high-throughput nature of our assay combined with automated tracking, we can easily test a large number of flies and song parameters, including those only rarely produced by conspecifics but to which animals might be sensitive (Coen et al. 2014; Aranha et al., 2017; Bussell et al., 2014; Crossley et al., 1995; Eberl et al., 1997; Rybak et al., 2002a; 2002b; F. Von Schilcher, 1976; Yoon et al., 2013; Zhou et al., 2015). Using FLyTRAP, we systematically compared male and female locomotor tuning to 82 acoustic stimuli that span the features and timescales present in courtship song (see Supplemental Table 1). Typically, each stimulus was presented 23 times to 120 females and 120 males, generating >2500 responses per stimulus and sex (see Methods).

Previous studies that assayed either male-female copulation rates or male-male chaining often focused on behavioral selectivity for the interval between pulses in a pulse train (inter-pulse interval (IPI), Fig. 1A) (Bennet-Clark and Ewing, 1969; Rybak et al., 2002b; F. Von Schilcher, 1976; Zhou et al., 2015). We therefore started by examining behavioral tuning for IPI using the wild type strain NM91, whose acoustic response during courtship was previously characterized(Clemens et al., 2015; 2018a; Coen et al., 2016; 2014). Observed changes in speed were stimulus-locked, sex-specific and tuned to IPI (Fig. 1C). Varying stimulus intensity had minimal effect on pulse song responses (Fig. S1A, B). While females slowed down to pulse trains, males exhibited transient slowing at pulse train onset followed by a long-lasting acceleration. The transient component of the locomotor response was present for all stimuli (Fig. S1C, D, see S3A-C) and may correspond to an unspecific startle response to sound onset (Lehnert et al., 2013). The transient was also present in females but masked by the stimulus-dependent slowing that followed (Fig. 1C). Due to the briefness of the transient response, the integral change in speed following stimulus onset reflects mostly the speed during the sustained phase (Fig. S1C, D). For simplicity, we therefore used the full integral as an overall measure of behavioral tuning. We found that in FLyTRAP, female IPI tuning is a band-pass filter matched to the statistics of male song (Fig. 1D): the mode of the distribution of *Drosophila melanogaster* IPIs is centered between 30 and 50 ms and females decrease their speed most for the same IPI range, and less for shorter or longer IPIs. Males produced a similar band-pass tuning curve peaked at the same IPI range - but their locomotor response was opposite in sign (males accelerated, females decelerated). This is consistent with the results of other assays that have found band-pass tuning for IPI in both sexes (Bennet-Clark and Ewing, 1969; Rybak et al., 2002b; F. Von Schilcher, 1976; Zhou et al., 2015) and a sex-specific sign of locomotor responses (Crossley et al., 1995; F. Von Schilcher, 1976).

Interestingly, we found the behavioral tuning for IPI in seven additional wild type strains to still be sex-specific but different from strain NM91 (Fig. S2A) – these strains showed reduced tuning for IPI, even though these same wild type strains – including NM91 – display similar responses to song features in a more natural courtship assay (Clemens et al., 2017; 2015; Coen et al., 2016; 2014). This indicates that these other strains may require additional cues (e.g., pheromones or visual cues) not present in FLyTRAP to fully express their preference for conspecific song features. For all subsequent analyses of locomotor tuning in FLyTRAP, we therefore chose the NM91 strain since i) acoustic responses during courtship were previously characterized (Clemens et al., 2018a; 2015; Coen et al., 2016; 2014), ii) it produced responses to song that were similar to the genetic background used for calcium imaging experiments (Fig. S2B-E), and iii) produced song responses that were consistent with those found using other assays (Crossley et al., 1995; Li et al., 2018; F. Von Schilcher, 1976; Yoon et al., 2013) – for example showing slowing to pulse song in females versus acceleration to pulse song in males.

We next systematically varied parameters that characterize pulse song to cover (and extend beyond) the distribution of each parameter within *D. melanogaster* male song (just as we did for IPI) (see Fig. S3). We examined behavioral tuning in both sexes for parameters that varied on timescales of milliseconds (carrier frequency, pulse duration and IPI) to seconds (pulse train duration) (Fig. 1A). We found that male and female tuning curves are of opposite sign but similar shape for all pulse song features tested across time scales (Fig. 2A, B, see Fig. S3A-C for speed traces, S3D-F), and that the behavioral tuning for pulse parameters often overlapped the distribution found in natural song (Fig. 2C). While the behavioral tuning curves for all pulse song features on short time scales are band-pass with a well-defined peak, we found that tuning for pulse train duration was monotonous: both females and males increase their locomotor response with increasing pulse train duration up to four seconds (Fig. 2A, B). During natural courtship, pulse trains longer than four seconds are rarely produced (Coen et al., 2014) – these stimuli thus correspond to “supernormal“ stimuli which drive strong behavioral response probably due to integration over long time scales (Tinbergen, 1989). Males also produce two distinct types of pulses (Clemens et al., 2018a) – we find that while females appear to be broadly tuned for both types of pulses in the FLyTRAP assay, males respond preferentially to higher frequency pulses (Fig. 2A, B). Finally, we found that both males and females are more selective for the pulse duration versus the pulse pause, the two components of the IPI (Fig. S3D-F) – this is in contrast to other insects that produce and process song pulses (e.g. crickets, grasshoppers, katydids), and that are preferentially tuned to pulse pause, pulse period or pulse train duty cycle (Hennig et al., 2014; Ronacher et al., 2014).

**Figure 2.**
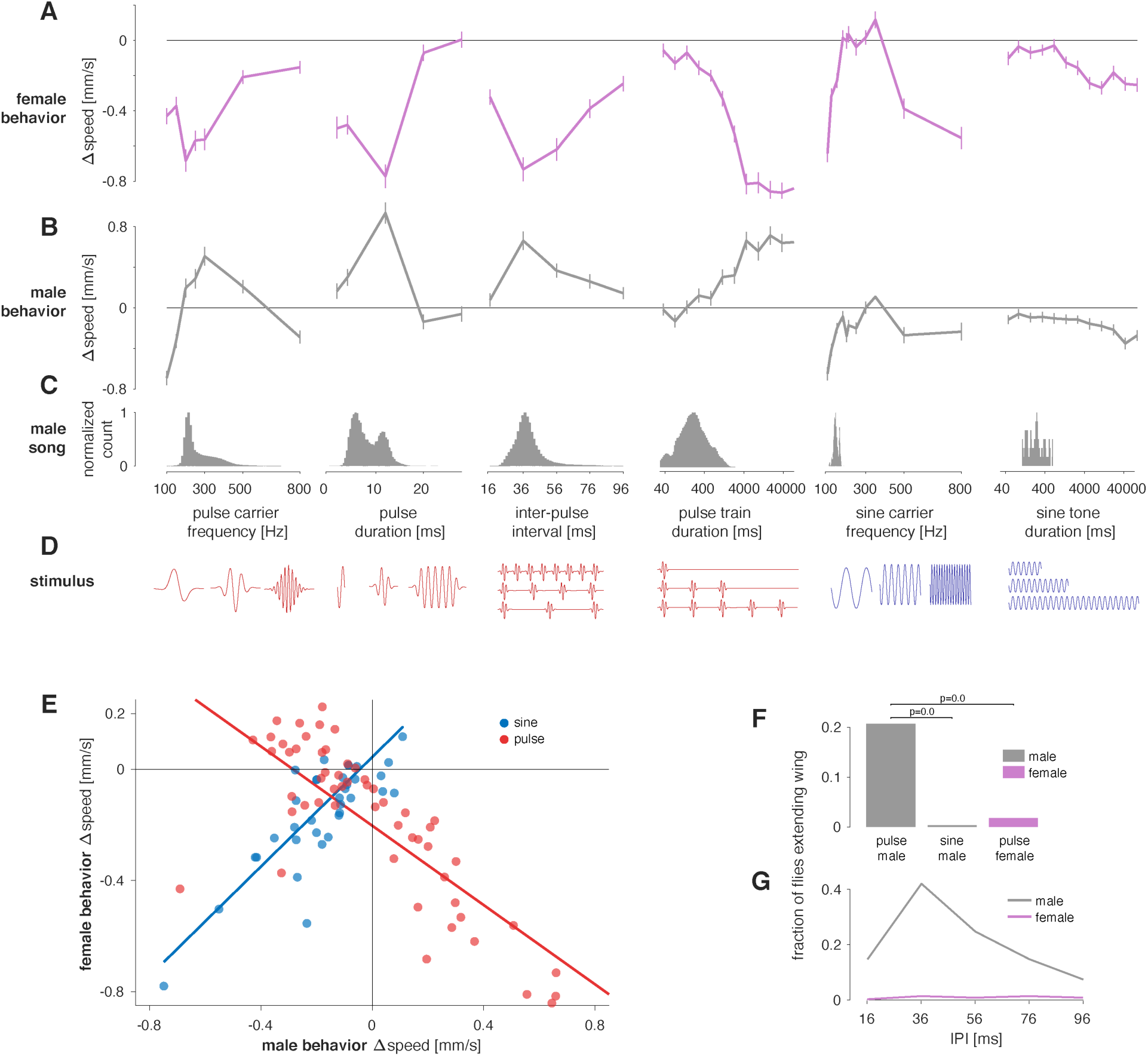
Responses to song playback are sex-specific and tuned for multiple features of pulse and sine song. **A, B** Locomotor tuning curves for females (A, magenta) and males (B, grey) for 6 different features of pulse and sine song. Lines and error bars correspond to the mean±s.e.m. across flies (see Table S1 for a description of all stimuli and N flies). **C** Distribution of the six different song features tested in A, B in the natural courtship song of *Drosophila melanogaster* males (data from 47 males of NM91 wild type strain totaling 82643 pulses and 51 minutes of sine song from 5269 song bouts). Histograms are normalized to a maximum of 1.0. **D** Pictograms (not to scale) illustrating each song feature examined in A-C. Pulse and sine song features are marked red and blue, respectively. **E** Changes in speed for males and females for all pulse (red) and sine (blue) stimuli tested (data from A, B, S3, see Table S1). Responses to sine stimuli are strongly and positively correlated between sexes (r=0.89, p=6×10^-8^). Pulse responses are also strongly but negatively correlated (r=-0.63, p=5×10^-10^). Blue and red lines correspond to linear fits to the responses to sine and pulse song, respectively. **F** Fraction of trials for which male and female flies extended their wings during the playback of pulse song (five different IPIs as in 1C, D) and sine song (150Hz, quantified only for males). Solitary males (grey) frequently extend their wings in response to pulse but not to sine song. Solitary females (magenta) do not extend wings for pulse song. See also Movie S2. **G** Fraction of trials that evoke wing extension in males (grey) and females (magenta) as a function of IPI. In males, wing extension and locomotor behavior (Figure 1D) exhibit strikingly similar tuning with a peak at the conspecific IPI. Females almost never extend their wing for any IPI. All behavioral data from the wild type strain NM91. All correlation values are Spearman’s rank correlation. See also Figures S2, S3 and Movie S2.

We next tested locomotor tuning for the parameters that characterize sine stimuli – carrier frequency and the duration of sine trains (Fig. 1A). Both males and females slow for sine tones of different frequencies, with very low and very high frequencies eliciting the strongest responses (Fig. 2A, B and Fig. S3A-C). Notably, the frequencies inducing the strongest slowing (100 Hz) are not typically produced by males (Fig. 2C). As for sine train duration tuning, we observed sustained responses that increased with duration and saturated only weakly, possibly because of the weak response magnitude.

Pulse and sine song usually co-occur within a single bout but it is not known why males produce two different modes (although females respond to both during natural courtship (Clemens et al., 2017; Coen et al., 2014)). One possibility is that one mode exerts a priming effect on the other (F. V. Schilcher, 1976). To test interactions between the two song modes, we presented sequences in which a 2-second pulse train was followed by a 2-second sine tone or in which a sine tone was followed by a pulse train and compared the responses for these sequences to the responses to an individual pulse train or sine tone (Fig. S3G). The responses are well explained by a linear combination of the responses to individual sine or pulse trains. Deviations from linearity occur due to sound onset responses, but otherwise responses do not strongly depend on the order of presentation in a bout (see also (Talyn and Dowse, 2004)). This suggests that these stimuli are processed in independent pathways.

To summarize, we compared behavioral responses in males and females from the strain NM91 for all features that define the courtship song. We found that male and female speed changes were strongly correlated for both song modes but that the sign of the correlation was negative for pulse stimuli and positive for sine stimuli (Fig. 2E). The opposite sign of the correlations along with the independence of responses to sine and pulse stimuli (Fig. S3G) indicates that sine and pulse song are processed by different circuits. The large magnitude of the correlations implies that feature tuning of the behavioral responses is similar between sexes and suggests that detector neurons for each song mode should be shared between sexes.

### Hearing pulse song drives wing extension in males, but not in females

Another sex-specific aspect of song responses is courtship: playback of conspecific song induces courtship-like behavior in males – this can even be directed towards other males, leading to the male chaining response, in which males follow other males, chasing and extending their wings (Eberl et al., 1997; F. Von Schilcher, 1976; Yoon et al., 2013). In our single-fly assay, males lack a target for courtship and the song-induced arousal likely manifests as an increase in speed. Since FLyTRAP does not permit simultaneous recording of fly acoustic signals during playback, we quantified wing extension as a proxy for singing, and examined whether song playback alone drives singing in solitary males. We found that solitary males extend their wings in response to pulse song stimuli specifically (Fig. 2F, G, Movie S2). This behavior is tuned for the inter-pulse interval (similar to the locomotor response, Fig. 1D) – the conspecific IPI of 36 ms drives the most wing extension, and shorter and longer IPIs evoke fewer wing extensions. By contrast, conspecific sine song (150 Hz) does not induce wing extension (Fig. 2F) (see also (Eberl et al., 1997; Yoon et al., 2013)). We also found that playback of pulse does not elicit wing extension in females, even though females have been shown to possess functional circuitry for singing (Clyne and Miesenböck, 2008; Rezával et al., 2016) – wing extension in response to pulse song is thus sex-specific.

These results are consistent with those for locomotor tuning: pulse song, but not sine song, generates sex-specific differences in the behavior of the strain NM91. The identical tuning of the two behavioral responses in males (locomotion (Fig. 1C) and song production (Fig. 2G)) suggests that the behavioral responses are driven by a common circuit.

### *Drosophila* male and female brains share pulse song detector neurons

Our systematic exploration of song stimulus space using the FLyTRAP assay revealed behavioral tuning for song parameters across temporal scales (from the carrier frequencies of sine and pulse lasting milliseconds to the duration of sine and pulse trains lasting seconds). We next searched for neurons with tuning across temporal scales that detect either the pulse or sine mode of courtship song. We focused on neurons expressing the Doublesex (Dsx) transcription factor that regulates sexual dimorphism in cell number and neuronal morphology between males and females. In the central brain there are ∼70 Dsx+ neurons per hemisphere in females and ∼140 Dsx+ neurons per hemisphere in males (Kimura et al., 2015; Rideout et al., 2010). Previous studies found calcium responses to both song-like stimuli and pheromones in Dsx+ neuron projections in females (Zhou et al., 2014) and tuning for the inter-pulse interval in males (Zhou et al., 2015). In addition, silencing subsets of Dsx+ neurons in females affected receptivity (Zhou et al., 2014). These data suggest that Dsx+ neurons could serve as the common pulse song detectors in males and females. To test this possibility, we recorded auditory responses in Dsx+ neurons and examined tuning for song features across timescales, in both males and females, to compare with our behavioral results.

We imaged neural activity using the calcium sensor GCaMP6m (Chen et al., 2013) expressed only in Dsx+ neurons. While we found no auditory response in the superior medial protocerebrum (SMP), we did find responses in the lateral junction (LJ) (Cachero et al., 2010; Ito et al., 2014; Yu et al., 2010), a site of convergence for the majority of Dsx+ neuron projections (Fig. 3A, B, S5B, C, Movie S3, S4). We found that male and female Dsx+ projections in the LJ are driven strongly by pulse but not by sine stimuli (Fig. 3C), confirming previous results (Zhou et al., 2014). While males produce weaker responses to auditory stimuli compared with females (Fig. 3C), the selectivity of Dsx+ LJ responses is highly correlated between sexes – stimuli that evoked the strongest responses in females also evoked the strongest responses in males (Fig. 3D).

**Figure 3.**
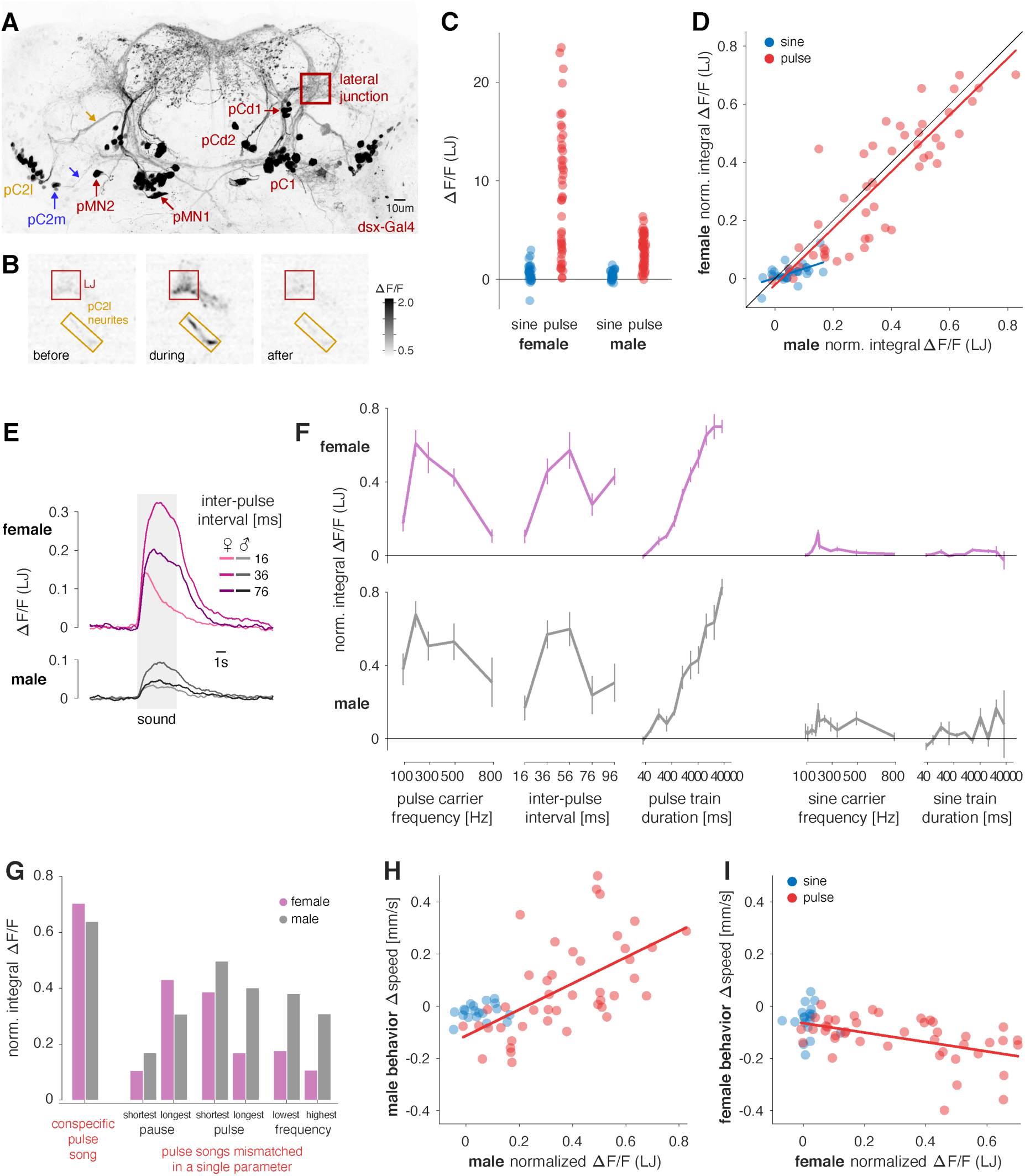
Neuronal tuning of Dsx+ neurons in the LJ matches behavioral tuning for pulse stimuli in males and females. **A** Anatomy of Dsx+ neurons in the female brain. Max z-projection of a confocal stack of a fly brain in which all Dsx+ are labeled with GFP. 5/8 cell types (pC1, pC2l (yellow), pC2m (blue), pMN1, pMN2) project to the lateral junction (LJ), while 3 cell types (pCd1, pCd2, aDN (movie S11)) do not. Yellow and blue arrows point to the neurites that connect pC2l and pC2m to the LJ. See also Fig. S5B, C. **B** Grayscale image (see color bar) of calcium responses (ΔF/F) to a pulse train (IPI 36ms) in an ROI centered around the LJ (red) and the pC2l neurites (yellow) in a female. Shown are snapshots of the recording at three different time points relative to stimulus onset - before (T=-10s), during (T=1.2s), and after (T=20s) the stimulus. Flies express GCaMP6m in all Dsx+ cells. Conspecific pulse song elicits strong increases in fluorescence in the LJ and the pC2 neurites. **C** LJ responses to sine (blue) and pulses (red) stimuli in females (left) and males (right). Individual dots correspond to integral ΔF/F responses for individual stimuli averaged over the 3-12 individuals tested for each stimulus. Many pulse stimuli evoke much stronger responses than the most effective sine stimulus (p=8×10^-11^ for females and p=2×10^-11^ for males, two-sided rank sum comparison of sine and pulse responses). **D** Comparison of male and female LJ responses to sine (blue) and pulse (red) stimuli. Responses to both song modes are correlated strongly for pulse (r=0.85, p=1×10^-14^) and moderately for sine (r=0.48, p=0.007) stimuli. Individual dots correspond to the integral ΔF/F for individual stimuli averaged across animals. Before averaging, the responses of each animal were normalized to compensate for inter-individual differences in calcium levels (see methods for details). **E** Fluorescence traces from the LJ in females (top, magenta) and males (bottom, grey) for pulse trains with three different IPIs (see legend, average over 6 individuals for each sex). In both sexes, the LJ responds most strongly to the conspecific IPI of 36ms (Fig. 1D). Responses are much weaker for shorter (16ms) and longer (76ms) IPIs. Calcium responses in the LJ are smaller in males than in females (compare C). See also Supp. Movie S3, S4. **F** Tuning curves of calcium responses in the female (magenta) and the male (gray) LJ for features of pulse and sine song (compare to behavioral tuning in Fig. 2A, B). Lines and error bars correspond to the mean±s.e.m. across flies. Integral ΔF/F normalized as in D. **G** pC2 calcium responses to the conspecific pulse song (left), pulse song stimuli with a mismatch in a single feature (right) in males (grey) and females (magenta). A single mismatch reduces neuronal responses by at least 20% and up to 80%, indicating the high, multi-feature selectivity of pC2 in both sexes. The conspecific pulse song is shown as a reference (pulse duration 12ms, pulse pause 24ms, pulse carrier frequency 250Hz, 112 pulses). Mismatch stimuli differed only in a single parameter from the reference (shortest pause: 4 ms, longest pause: 84 ms; shortest pulse: 4ms, longest pulse: 60ms, lowest frequency: 100Hz, highest frequency: 800Hz). **H, I** Comparison of behavioral and neuronal tuning in males (H) and females (I). Behavioral and neuronal data from flies of the same genotype (Dsx/GCaMP). We obtained similar results when comparing the neuronal responses to behavioral data from wild type flies (NM91, Fig. S4H-I). Dots correspond to the Δspeed and the normalized integral ΔF/F averaged over individuals, lines indicate linear fits. In males (H), behavioral and neuronal responses are *positively* correlated for pulse (red, r=0.61, p=1×10^-5^) but not for sine stimuli (blue, r=0.17, p=0.49). In females (I), behavioral and neuronal responses are *negatively* correlated for pulse (red, r=-0.53, p=3×10^-4^ but not for sine stimuli (blue, r=0.28, p=0.25). All Δspeed and ΔF/F values are from Dsx/GCaMP flies. Panel K additionally shows behavioral data from the wild type strain NM91. All correlation values are Spearman’s rank correlation. See also Figure S4, and Movie S3 and S4. Figure 4

When examining neuronal tuning curves, we found a good match between Dsx+ LJ responses and the magnitude of changes in speed across all timescales of pulse song in both sexes (Fig. 3E, F). Results were similar whether we used the integral ΔF/F or the peak ΔF/F to quantify tuning (Fig. S4F, G). For example, the Dsx+ LJ tuning curve for IPI is similar in females and males with the strongest responses at 36 ms, matching the behavioral tuning curves (compare with Fig. 2A-B and S2B-C). On longer timescales, LJ tuning curves also match behavioral tuning curves for pulse train duration, with the integral calcium increasing with train duration in both sexes, similar to the behavioral response. Overall, LJ responses are highly selective for multiple features found in conspecific pulse song: Dsx+ LJ responses were strongest for stimuli with a carrier frequency of 250 Hz, an inter-pulse interval of 36 ms, and a pulse duration of 16 ms (Fig. S4C-E). A match in only two of these three features was not sufficient to maximally drive Dsx+ LJ projections (Fig. 3G) – a mismatch in a single pulse song feature reduced calcium responses between 20% and 80%. Sine stimuli have lower carrier frequencies, long durations, and no pauses (they are by definition continuous) – which explains the weak responses of Dsx+ LJ neurons to all sine stimuli (Fig. 3C). Likewise, broadband noise also lacks the correct pattern of amplitude modulations and accordingly does not strongly drive the Dsx+ LJ neurons (see also Fig. 4D). Given this high selectivity for the features defining conspecific pulse song, it is unlikely that the LJ would be driven by other naturally occurring stimuli. For instance, wind stimuli typically contain lower frequencies (Nagel and Wilson, 2011) and lack the periodical pattern of pulse trains required to strongly drive Dsx+ LJ neurons, and aggression song differs from courtship pulse song in carrier frequency and in IPI (Versteven et al., 2017).

**Figure 4.**
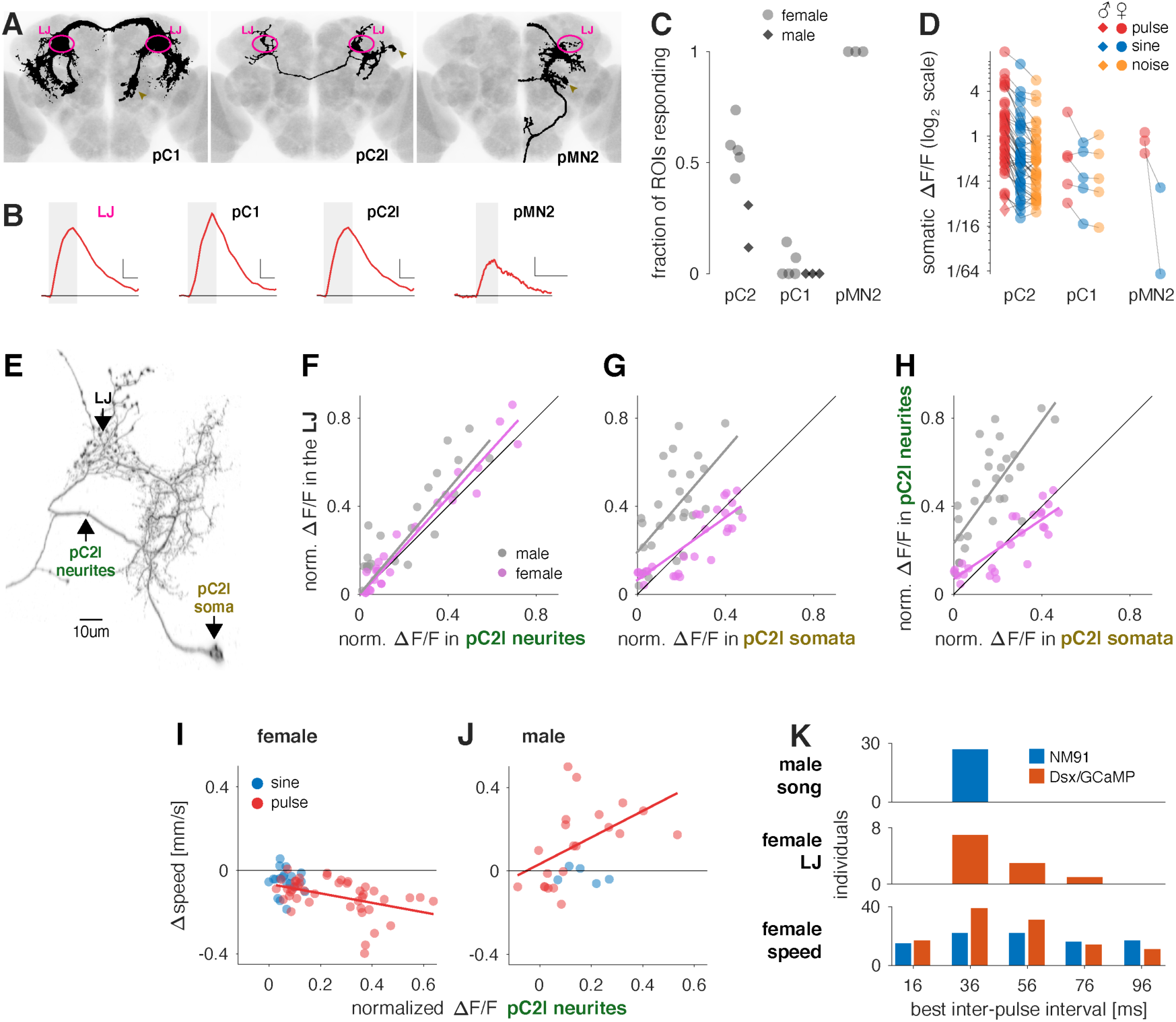
pC2 neurons are pulse song detectors common to both sexes. **A** Individual Dsx+ neuron types (black) with somas in the female central brain in which we detected calcium responses for pulse or sine song, registered to a common template brain (gray) (see Methods for details). Of the 8 Dsx+ cell types in the central brain, pC2l, pC2m, the single female-only neuron pMN2 and a small number of pC1 neurons (and only in some individuals) respond to courtship sounds. The lateral junction (LJ) is marked in magenta and somata are marked with golden arrow heads. See also Supp. Movie S10, S11. **B** Example somatic fluorescence traces from single somata of the pC1, pC2, and pMN2 cells in response to pulse trains (IPI=36ms, single trial responses). Fluorescence trace from the LJ (magenta) shown for comparison. The gray box marks the duration of the sound stimulus. In each panel. Horizontal and vertical scale bars correspond to 6 seconds and 0.25 ΔF/F, respectively. Horizontal black line marks ΔF/F=0. **C** Fraction of cells in Dsx+ clusters with detectable somatic calcium responses to pulse or sine song (females, light grey dots; males, dark grey squares). Complete clusters were imaged using volumetric scan for pC1, pC2 and single plane scans for pMN2. We did not distinguish between pC2l/m, since in most flies both groups are spatially intermingled at the level of cell bodies. Note that all flies included showed calcium responses to sound in the LJ, even when we did not detect responses in specific somata. **D** Peak somatic ΔF/F for pulse (red, 36ms IPI), sine (blue, 150Hz), and noise (orange, 100-900Hz). Lines connect responses recorded in the same animal. Note that responses are plotted on a log scale – the average of the ratio between sine and pulse for all cells is ∼2.6. 36/38 pC2, 4/5 pC1 and 2/2 pMN2 prefer pulse over sine. See also Supp. Movie S5, S6. **E** High resolution confocal scan of a single pC2l neuron (obtained via a stochastic labelling technique, see Methods for details). Only the side ipsilateral to the cell body is shown. The neurites in the lateral junction appear varicose, indicating that they contain pre-synaptic sites. **F** Normalized integral ΔF/F values recorded simultaneously in the LJ and the neurites that connect the LJ with the somata of pC2l (and no other Dsx+ cell type) are highly correlated in females (magenta, r=0.99, p=1×10^-71^, N=10-24 flies/stimulus) and males (grey, r=0.75, p=4×10^-13^, N=1-6 flies/stimulus). Each point corresponds to an individual stimulus (pulse or sine) averaged over flies. The high correlation indicates that calcium responses in the LJ reflect responses in pC2l neurons. Magenta and gray lines in F-H correspond to a least-squares fit to the individual data points. **G** Normalized integral ΔF/F recorded first in the LJ and then in single pC2l somata in the same fly are highly correlated in both sexes (females: r=0.86, p=8×10^-10^, N=8 flies/stimulus, males: r=0.73, p=4×10^-6^, N=1 fly/stimulus), demonstrating that calcium responses in the LJ represent the responses of individual pC2l cells, with some variability across individual cells and animals. **H** Normalized integral ΔF/F responses from the pC2l neurites and from single pC2l somata in different flies are highly correlated in both sexes (females: r=0.89, p=2×10^-11^, N=8 flies/stimulus, males: r=0.79, p=1×10^-7^, N=1 fly/stimulus). The pC2l neurites reflect the average activity of individual pC2l neurons, with some variability across individual cells and animals. **I, J** Comparison of calcium responses in the pC2l neurites and male (I) or female (J) speed for the same stimuli. Calcium and speed data come from flies of the same genotype (Dsx/GCaMP). Similar results were obtained when using speed data from wild type flies (NM91) instead (Fig. S5G-H). pC2l and behavioral responses are highly correlated for pulse with a sex-specific sign (female (I): pulse: r=-0.49, p=1×10^-3^, sine: r=-0.09, p=0.73; male (J): pulse: r=0.70, p=5×10^-4^, sine: r=-0.20, p=0.78), just as for the LJ (compare Fig. 3I). The match between neuronal and behavioral tuning for pulse song indicates that pC2l neurons detect the pulse song. Each point corresponds to an individual stimulus (Δspeed: N∼100 flies per stimulus, ΔF/F: N=10-24 female and 1-6 male flies/stimulus). **K** Comparison across individuals of most frequent IPIs in male song (n= 75528 pulses from 27 males) and preferred IPIs in the female LJ (integral ΔF/F; n = 11 females) and behavior (Δspeed; n = 112 females NM91 and 92 females Dsx/GCaMP). Song and speed are shown for NM91 (blue), LJ and speed are shown for Dsx/GCaMP (orange). While all males produce songs with IPIs around 36ms, female neuronal and behavioral tuning for IPI is much more variable (standard deviations: 2.4ms for male song, 14ms for female ΔF/F (for integral ΔF/F (shown), 7ms for peak ΔF/F), 23 and 27ms for the speed of NM91 and Dsx/GCaMP females, respectively). Notably, variability in female speed is larger than in the female LJ, indicating that pathways parallel to or downstream of the LJ contribute to the behavior. All Δspeed and ΔF/F values are from flies expressing GCaMP6m under the control of Dsx-Gal4. All correlation values are Spearman’s rank correlation. See also Figure S5 and Movie S5 and S6.

To more directly show that Dsx+ LJ neurons are shared pulse song detectors, we matched neuronal and behavioral responses for the same stimuli. Given the observed strain-dependence of behavioral responses in FLyTRAP (Fig. S2A), we compared behavioral and neuronal tuning within the same genotype (Dsx/GCaMP). In FLyTRAP, Dsx/GCaMP flies produced weaker behavioral responses, but nonetheless, their tuning for song features was matched to that of wild type strain NM91 (Fig. S2B-E). Male neural and behavioral tuning for pulse stimuli are positively correlated (r=0.61, p=1×10^-5^) – high Dsx+ LJ neuron activity correlates with the most acceleration (Fig. 3H). Female neural and behavioral tuning for pulse stimuli are negatively correlated (r=-0.53, p=3×10^-4^) – high Dsx+ LJ neuron activity correlates with the most slowing (Fig. 3I). This is consistent with these neurons controlling the magnitude, but not the direction of speed changes. We observed no statistically significant correlation for sine stimuli (male: r=0.17, p=0.49; female: r=0.28, p=0.25), as Dsx+ LJ responses to sine stimuli were weak. We obtained a similar pattern of correlations when using behavioral data from the wild type strain NM91 for comparison instead (Fig. S4H-I). Note that Dsx+ LJ activity only accounts for roughly 1/3 of the variability in behavioral responses for pulse song. This suggests that the behavior is driven and modulated by additional pathways outside of the Dsx+ neurons in the LJ or that most of the variability arises in downstream circuits. Nonetheless, Dsx+ neurons that innervate the LJ have tuning properties expected for pulse song detectors – they prefer pulse over sine stimuli, are similarly tuned in males and females, and their feature tuning matches the behavioral tuning for all pulse, but not sine, stimuli across timescales.

### Dsx+ pC2 neurons are tuned like the LJ and to conspecific pulse song

The Dsx+ neurons of the central brain form a morphologically heterogeneous population with several distinct, anatomical clusters many of which project to the LJ (Kimura et al., 2015; Rideout et al., 2010; Zhou et al., 2015; 2014) (Fig. 3A). Previous studies that examined auditory responses in Dsx+ neurons (Zhou et al., 2015; 2014) did not resolve which subtype carried the response. Using a stochastic labeling approach (Nern et al., 2015), we confirmed that five out of eight Dsx+ cell types in the female brain have projections into the LJ (Kimura et al., 2015): pC1, pC2l/m, pMN1, and pMN2, but not pCd1/2 and aDN (Fig. 4A and Fig. S5D, E). We next imaged calcium responses to pulse and sine stimuli in the somas of all five Dsx+ cell types that innervate the LJ and found that a subset of neurons in the pC1 and pC2 clusters possess auditory responses, in addition to cell type pMN2 (a VNC-projecting female-specific neuron (Kimura et al., 2015) comprising only one cell body per hemisphere) (Fig. 4B, C, Movie S5, S6). All responsive cells preferred pulse over sine or noise stimuli (Fig. 4D). We did not observe auditory responses in pMN1 neurons (not shown), although we cannot rule out that this neuron class has responses that are below the level of detection by the Calcium indicator GCaMP6m.

The pC1 cluster – which was previously considered the only Dsx+ auditory neuron in the LJ (Zhou et al., 2015; 2014) – contained very few somas with calcium responses to sound (2-3 cells in the female brain; none in the male brain) (Fig. 4C). By contrast, we found ∼15 auditory neurons in the pC2 cluster in each animal (this number is an underestimate since somas overlap; see Methods). While pC1 and pMN2 likely contribute to the LJ responses, they contain few auditory-responsive neurons and/or are present only in females. We therefore focused on pC2 as the putative pulse song detector common to both sexes.

Although there are more pC2 neurons in males versus females (∼67 vs. ∼26, (Kimura et al., 2015)) the number of auditory neurons is similar in both sexes (∼15). pC2 neurons can be subdivided into a lateral and a medial type, termed pC2l and pC2m (Robinett et al., 2010), and each type projects to the lateral junction via a distinct bundle of neurites (see Fig. 3A, S5B, C). Most auditory neurons were lateral in the pC2 cluster in both sexes (Fig. S5A), and pC2l neurites produced strong auditory responses. However, we did observe auditory responses from some pC2m neurons indicating that auditory activity is not exclusive to pC2l (Fig. S5A). While tuning differed slightly between individual pC2 neurons, no single cell was specialized to detect specific features of the pulse song (Fig. S5F) and responses of single cells and the LJ were highly correlated in both sexes (Fig. 4F-H). From this we conclude that LJ responses reflect the tuning of pC2l neurons. Importantly, the tuning of the pC2l neurites in the LJ matches the behavioral tuning in both sexes (Fig. 4I-J and Fig. S5G-H), indicating that pC2l neurons are selective for conspecific pulse song.

### Circuits parallel to or downstream of pC2l strongly contribute to behavioral variability

The match between behavioral and pC2 tuning suggests that pC2 contributes to the sex-specific responses to song. If locomotor responses were driven mainly by pC2, then the variability in pC2 tuning across animals would explain most of the variability in behavioral responses across animals. On the other hand, if locomotor responses were controlled by parallel pathways or by circuits downstream of pC2, then the variability in pC2 would be much lower than the behavioral variability across animals. We compared individual variability of male song, female pC2 neural responses, and female locomotor responses (Fig. 4K) and focused on the pulse song feature for which we had the most data – inter-pulse interval (IPI). We found a steady increase in variability from song to brain to behavior. Song is consistent across individuals – the IPI distribution for each male peaks between 36 and 56 ms. pC2 (LJ) responses are more variable than song but still relatively consistent across animals – pC2 in 7/11 females responds most strongly between 36 and 76 ms. By contrast, behavioral responses are highly variable – only half of the flies slow most strongly to IPIs between 36 and 76 ms. Variability at the level of locomotor responses increases for other song features, too (data not shown). Overall, this suggests that locomotor responses in FLyTRAP are strongly affected by pathways parallel or downstream of pC2. This must be considered when interpreting experiments that test the role of pC2 in driving behavioral responses to song.

### Activation and inactivation of pC2l neurons affects sex-specific behaviors

Given that pC2l neurons are tuned to the conspecific pulse song, we expected that their activation could also contribute to the sex-specific behaviors observed for pulse song – changes in locomotion and singing that are distinct between males and females. To test this hypothesis, we used a driver line (Rezával et al., 2016; Zhou et al., 2014)) that labels 11/22 female and 22/36 male pC2l neurons, in addition to 5-6 pCd neurons, but no pC2m or pC1 neurons (Fig. S6A). At least 5 of the pC2l cells in this driver line responded to song (Fig. S6B), which corresponds to ∼1/3 of the auditory pC2l neurons. We expressed CsChrimson, a red-shifted channelrhodopsin (Klapoetke et al., 2014), in these neurons and optogenetically activated them in both males and females.

We first recorded behavior in a chamber tiled with microphones (Coen et al., 2014) to test whether pC2 activation was sufficient to induce singing, as we previously showed that pulse song playback alone drives wing extension in males (Fig. 2F, G). Upon red light activation, males produced pulse song, while sine song was produced transiently after stimulus offset (Fig. 5A, Movie S7), and the amount of pulse song produced scaled with the strength of activation (Fig. 5B-C). The evoked pulse and sine songs were virtually indistinguishable from natural song (Fig. S6C, D). In *Drosophila*, retinal (the channelrhodopsin cofactor) must be supplied via feeding, and red light stimulation drove singing significantly more in males fed with retinal versus those fed regular food (Fig. S6E). Activation of a control line that only labels pCd neurons (Zhou et al., 2014) did not drive singing (Fig. S6E), implying that song production results from the activation of the pC2 neurons in our driver. Importantly, we never observed song production upon pC2 activation in females (Fig. 5D-E) – pC2 neurons thus drive song in a sex-specific manner. These results also establish pC2 neurons as serving a dual sensory and motor role: they respond selectively to the pulse song (Fig. 3C, F) and also bias the song pathway towards producing the same song mode (Fig. 5A, B).

**Figure 5.**
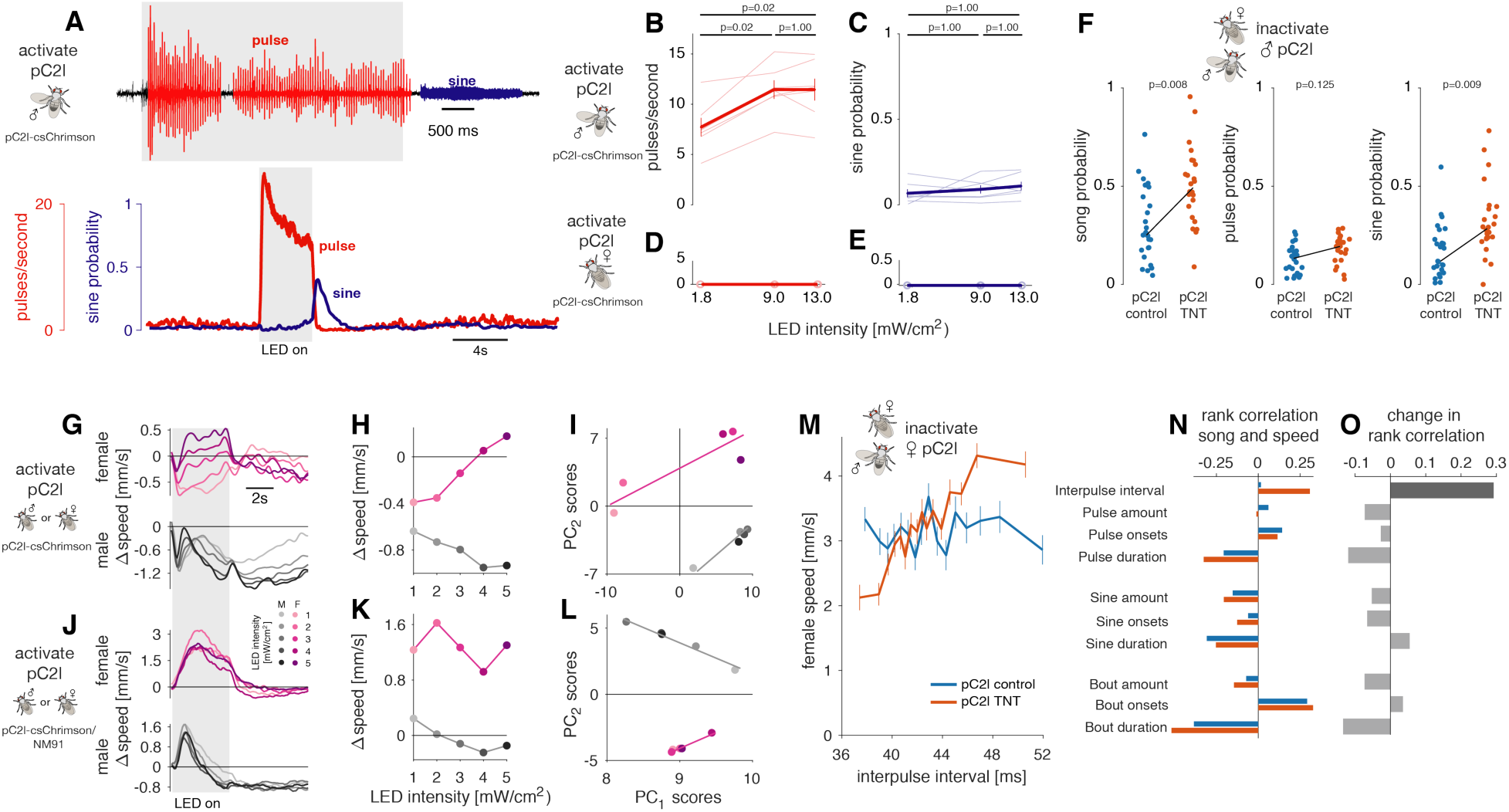
Testing the necessity and sufficiency of pC2 neurons for song and locomotor behaviors. **A** Song evoked in males by optogenetic activation (627nm LEDs, intensity 13 mW/cm^2^) of a driver line that labels pC2l and pCd neurons (R42B01∩Dsx, refered to as pC2l-csChrimson). Top trace shows a song recording marking pulse and sine song in red and blue, respectively. The grey area indicates the duration (4 seconds) of optogenetic activation. Pulse song is evoked during activation while sine song occurs immediately following activation. Bottom plots show pulse rate (red) and sine song probability (blue) averaged over 7 flies (18 stimulation epochs per animals). See also Movie S7. Activation of pC2l using a different genotype (pC2l-csChrimson/NM91) has similar effects (Fig. S6I, J) **B, C** Average pulse rate (B) and sine song probability (C) evoked in the 6 seconds following LED light onset (LED duration is 4 seconds). Dose-response curves for individuals are shown as thin lines, population averages (mean±s.e.m.) are shown as thick lines with error bars. P-values result from two-sided sign tests and are adjusted for multiple comparisons using Bonferroni’s method. Same data as in A. See also Supp. Movie S7, S8. **D, E** Same as B, C but for females (N=3 flies). Activation of pC2l (and pCd) in the female does not evoke song – pC2l activation drives singing in a sex-specific manner. **F** Song of males courting wild type NM91 females. pC2l synaptic output in the males was inhibited using TNT via the R42B01∩Dsx driver. Dots correspond to the amount of all song (left), pulse song (middle), and sine song (right) (pC2l TNT (N=24) – orange, pC2l control (N=25) – blue). Black lines connect the means of the two genotypes. P-values show the outcome of a two-sided rank sum test. Inhibiting pC2l output leads to more overall singing and sine song, but not to more pulse song, indicating that pC2l biases singing towards pulse song during courtship. Other song features are not affected (see Fig. S6F-H). **G, H** Optogenetic activation of R42B01∩Dsx using csChrimson (pC2l-csChrimson) evokes locomotor responses with sex-specific dynamics. Changes in speed (G) and tuning curves (H) were corrected for intrinsic light responses by subtracting the responses of control flies with the same genotype that were not fed retinal (see Fig. S6K). Females (top, magenta) slow for weak and speed for strong activation with multi-phasic dynamics as for sound (H, I, compare D). Males decrease their speed and responses outlast the optogenetic stimulus (bottom, grey). See also Supp. Movie S9. See S6K for N flies. The grey area indicates the duration of LED stimulation (4 seconds). **I** Principal component (PC) analysis of male and female locomotor speed traces (12s following stimulus LED or sound onset, traces taken from G). Shown are first and second principal component (PC) scores of females (magenta) and males (grey) for sound (squares) and optogenetic stimulation (circles). Lines correspond to least-square fits for each sex. Female and male responses to different LED occupy different areas in PC space, indicating that the locomotor dynamics are sex-specific. **J, K** Same as G, H but with a different genotype (pC2l-csChrimson/NM91 – see Methods for details). Females (top, magenta) speed throughout the stimulation (J) and for all LED intensities (K). Males (bottom, grey) first speed and then slow for all LED intensities. The evoked locomotor dynamics differ between genotypes (I) but are always sex-specific. **L** Same as I but with the pC2l-csChrimson/NM91 phenotype. Again, male and female locomotor responses are different, since they occupy different areas in PC space (compare panel I). **M** Locomotor tuning for IPI during natural courtship obtained from single females that were courted by a wild-type NM91 male. pC2l synaptic output in the females was inhibited using TNT using the R42B01∩Dsx driver. Lines and error bars correspond to the mean±s.e.m speed of N females per genotype tested (pC2l TNT– orange, pC2l control – blue, N=48 females for each genotype, see methods for details on how the tuning curves were computed). pC2l control females (blue) do not change their speed with IPI within the range commonly produced by males (r=0.02, p=0.59, compare Fig. 1D). pC2l TNT females (orange) accelerate for longer IPIs (r=0.31, p=3×10^-30^). **N** Rank correlation between female speed and different song features during natural courtship (pC2l control – blue, pC2l TNT – orange). **O** Difference between the rank correlations for control (blue) and pC2l TNT (orange) flies in N. pC2l inactivation specifically changes the correlation between female speed and IPI (dark gray, p=6×10^-8^). All other changes in correlation are much smaller and not significant (p>0.18). P-values were obtained by fitting an ANCOVA model (see methods for details) and were corrected for multiple comparisons using the Bonferroni method. All correlation values are Spearman’s rank correlation. See also Figure S6 and Movie S7.

We next tested whether inactivation of pC2 affected song production during courtship, by constitutively suppressing the synaptic output of pC2 (via expression of TNT (Sweeney et al., 1995)) in males courting wild type virgin females (see Methods). Males with a genetically silenced subset of pC2 neurons still sang, demonstrating that the driver did not label neurons that were required for singing or that other neurons can substitute for the missing activity of pC2 (Clemens et al., 2018a; Philipsborn et al., 2011). The fine structure of the song was wild-type like with normal IPIs, pulse shapes, and carrier frequencies (Fig. S6F, G) and copulation rates were normal (Fig. S6H). Surprisingly, pC2l-silenced males sang about twice as much as the controls, and this effect was largely driven by the production of more sine song (Fig. 5F). Given that pC2 activation yielded virtually no sine song during optogenetic stimulation (Fig. 5A, B), this suggests that pC2 inhibits sine song production during natural courtship and generally demonstrates that song production in *Drosophila* involves a complex control scheme (see also (Clemens et al., 2018a; Philipsborn et al., 2011; Shirangi et al., 2013)).

The genetic manipulations so far demonstrate that pC2 can drive behavior in a sex-specific manner – driving song only in males. FLyTRAP also revealed pC2 sex-specific locomotor responses. In the wild type strain NM91, these responses were similarly tuned but of opposite sign in both sexes (Fig. 1, 2). Although the behavioral tuning differed for other strains, the locomotor responses to song were still sex-specific (Fig. S2A). To test whether pC2 activation can produce sex-specific locomotor responses, we placed flies in the FLyTRAP assay and used red light for activation (instead of sound). Given the genotype dependence of the locomotor tuning, we expressed csChrimson in pC2 using two different genotypes. Both carried the same transgenes for expressing csChrimson in pC2 neurons, but one carried half of its chromosomes from the NM91 wild type strain – these genotypes are called “pC2l-csChrimson” and “pC2l-csChrimson/NM91” (see Methods). Both strains produced song upon optogenetic activation in males but not in females (Fig. 5A-E, S6I, J). In FLyTRAP, these strains produced different but nonetheless sex-specific locomotor responses for IPI stimuli (Fig. S2A), allowing us to test whether locomotor responses evoked by pC2 activation are robustly sex-specific despite genotype-specific locomotor tuning. To account for innate visual responses to the light stimulus, we subtracted the responses of normally fed flies from retinal fed flies (Fig. S6K, L).

For both strains, optogenetic activation of pC2 yielded sex-specific locomotor responses. For pC2l-csChrimson, we observed complex, multiphasic locomotor dynamics, with males tending to slow down and females tending to speed up with increasing optogenetic activation (Fig. 5G, H). For pC2l-csChrimson/NM91, we observed simpler, bi-phasic responses – females first sped up during activation and slowed down after, while males sped up for a short period after stimulation onset only (Fig. 5J). For this genotype, responses differed little across activation levels (Fig. 5K). Importantly, locomotor responses were sex-specific in both genotypes, which we confirmed using principal component analysis (PCA) of the speed traces of males and females (Fig. 5I, L). The first two principal components were sufficient to explain 80% and 99% of the variance in the speed traces, and the responses occupy non-overlapping regions in the principal component space. However, pC2 activation in neither strain reproduced the responses to pulse trains of varying IPI (for the same strain) in FlyTRAP (cf. Fig. S2A). This could be because optogenetic activation does not recapitulate brain dynamics evoked by song – either because the pC2 activation levels were not matched or because song activates multiple circuits that all affect the locomotor responses (Jazayeri and Afraz, 2017). While these issues have to be addressed to fully understand how the responses to playback of song are driven, the results show that pC2 is one of serval elements that contribute to the locomotor tuning for song.

Finally, we used the pC2l-csChrimson driver to constitutively suppress the synaptic output of pC2 (via expression of TNT (Sweeney et al., 1995)) in females and paired them with wild type virgin males (see Methods). We quantified female song responses as the correlation between different song features and female speed (Clemens et al., 2015; Coen et al., 2014) (Fig. 5M-O). Because male song is structured via sensory feedback cues from the female (Coen et al., 2014), silencing pC2 neurons in females could affect the content of male song – however, the statistics of male song were unchanged by the female manipulation (Fig. S6M, N). pC2 inactivation specifically affected the correlation between female speed and the pulse song IPI, which changed from ∼0 to +0.3 (Fig. 5M-O). While control – and wild type (Clemens et al., 2015) – females do not change their speed relative to the range of natural IPIs produced by conspecific males (Fig. 1), females with pC2 neurons silenced accelerate more with increasing IPI. pC2 neurons are therefore required for the normal response to pulse song. The remaining responses to pulse could be caused by pC2 neurons not silenced by our genetic driver or by other neurons tuned for longer IPIs (Zhou et al., 2015). While female locomotor responses to courtship song were affected by pC2 inactivation, copulation rates were not significantly reduced (Fig. S6O), consistent with previous studies (Zhou et al., 2014). In conjunction with the match between behavioral tuning and pC2 tuning, these results add to the evidence that pC2 neurons detect pulse song and play a critical role at the sensorimotor interface – they relay information about pulse song to sex-specific downstream circuits that control either singing or locomotion, and thereby contribute to acoustic communication behaviors.

### Auditory responses of pC2 are modulated by social experience

Many sexual behaviors change with social experience (Keleman et al., 2012; Li et al., 2018; Marlin et al., 2015; Remedios et al., 2017). This plasticity could be mediated by modulating the selectivity and the gain of the neurons that detect social cues or of the neurons that drive the behaviors. Social experience is also known to affect courtship behavior in Drosophila (Ellis and Kessler, 1975; Kohatsu and Yamamoto, 2015; F. Von Schilcher, 1976). In particular, a recent study has shown that group housing sharpens the IPI selectivity of the female mating decision and of the male chaining response, and that this effect is mediated by the exposure of song from other flies in the group (Li et al., 2018). However, we do not yet know which elements in the pathway from song to behavior are affected by social experience. Given that pC2 contributes to behavioral responses to song, we asked whether its activity is modulated by housing conditions. The behavioral results presented so far were obtained from group-housed flies so we also ran single-housed males or females to confirm that locomotor responses in FLyTRAP are modulated by social experience. We found that single-housed males responded with little selectivity to pulse trains with different IPIs (Fig. 6A). This is consistent with the previous study (Li et al., 2018), since group-housed males are exposed to the song of other males during rearing. That we can reproduce these results in a single-fly assay shows that acoustic cues are sufficient to express the effect – previous experiments had used multi-fly assays, leaving open the possibility of other cues being required – e.g. social experience could also affect the song selectivity by altering the pheromonal modulation of acoustic responses. By contrast, females do not sing to other females and accordingly, their locomotor responses are unaffected by the housing condition. Consistent with the behavior, calcium responses in pC2 (measured via the LJ) (Fig. 4F-H) do not change strongly with housing conditions in females but become more selective for IPI in group-housed males (Fig. 6B, C). Notably, sine song responses and responses to pulse trains with different durations are not affected by housing conditions (Fig. S7). This suggests that pC2 could mediate the effect of social experience on the behavioral responses to song.

**Figure 6.**
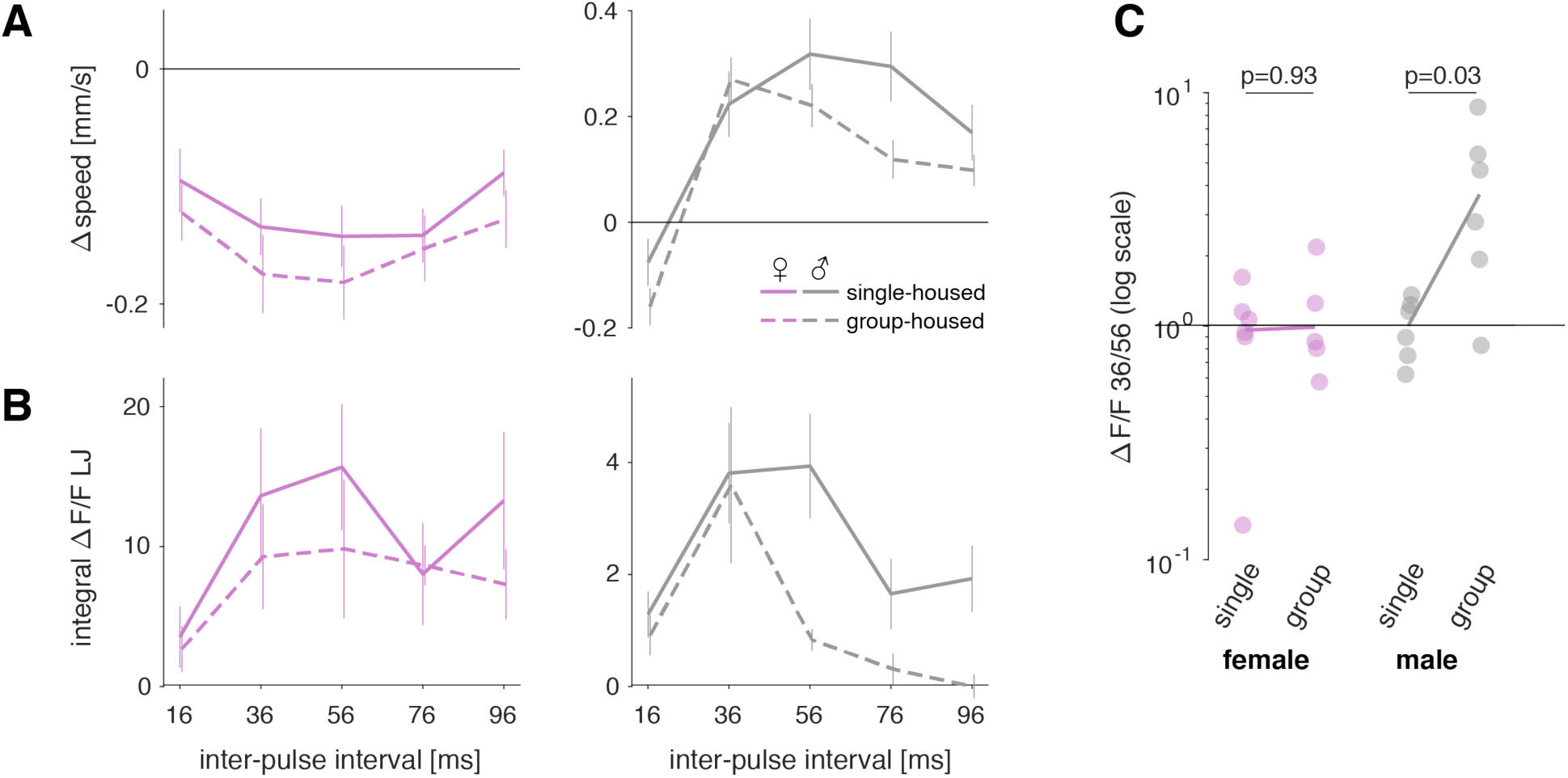
Behavorial and pC2 responses are similarly modulated by social experience. **A** Changes in speed for pulse trains measured using FLyTRAP with different IPIs in single-housed (solid line) or group-housed (dashed lines) female (left, magenta) and male flies (right, grey). Plots show mean ± s.e.m. across 92/116 group-housed and 137/71 single-housed female/male flies. Female IPI tuning is not strongly affected by housing conditions. By contrast, males change their speed more selectively when group-housed. **B** Calcium responses from the LJ for pulse trains with different IPIs in single-housed (solid line) or group-housed (dashed lines) female (left, magenta) and male flies (right, grey). Plots show mean ± s.e.m. across 5-6 female or male flies in each condition. In females, group housing only weakly suppresses LJ responses for some IPIs. By contrast, male LJ responses are selectively suppressed for long IPIs, which sharpens the IPI tuning. **C** Ratio of Calcium responses to 36 and 56 ms IPIs in single-housed or group-housed female (left, magenta) and male flies (right, grey). Individual dots correspond to individual flies, the solid lines connect the population average ratios. P-values were obtained from a two-sided rank sum test. All Δspeed and ΔF/F values are from flies expressing GCaMP6m under the control of Dsx-Gal4. See also Figure S7.

## Discussion

Using a quantitative behavioral assay, we characterized locomotor responses in both males and females to the features that define the *Drosophila melanogaster* courtship song. Males and females of the wild type strain NM91 showed similar tuning for pulse song stimuli, but nonetheless produced distinct responses (males accelerate while females decelerate; males sing while females do not) (Fig. 1, 2). For both males and females and across multiple timescales, tuning was matched to the distribution of each parameter in the male’s pulse song. We then identified Dsx+ pC2 neurons in the brain that respond selectively to all features of pulse song stimuli, and whose tuning is matched to behavioral tuning (Fig. 3, 4). The activation of pC2 neurons elicited sex-specific behavioral responses to pulse song (Fig. 5), and social experience sharpened both behavioral feature selectivity and pC2 tuning (Fig. 6). We thus conclude that Dsx+ pC2 neurons connect song detection with the execution of sex-specific behaviors.

### Matches between behavioral tuning and conspecific song

Behavioral selectivity for species-specific signals is thought to serve species separation. In FLyTRAP, locomotor tuning in NM91 and Dsx/GCaMP females overlaps with the conspecific song – these females slow to conspecific song (Fig. 2A) and do not change their speed or may even accelerate for deviant pulse parameters (Fig. S3A, E). However, the tuning for any single song feature is not sufficiently narrow to serve as an effective filter for conspecific song. For instance, NM91 and Dsx/GCaMP females also slow for IPIs produced by a sibling species *D. simulans* (50-65 ms) (Bennet-Clark and Ewing, 1969). However, *D. simulans* pulses would be rejected based on a mismatch in other song features – *D. simulans* pulses are too short and of too high frequency to be accepted by females (Clemens et al., 2017; Riabinina et al., 2011). Selectivity for multiple song features may thus enable species discrimination with relatively broad single-feature tuning (Amézquita et al., 2011). Males and females are exposed to additional non-acoustic cues during courtship that may further sharpen behavioral tuning. For instance, chemical cues prevent males from courting heterospecific females (Fan et al., 2013) and likely also contribute to female rejection (Billeter et al., 2009; Rybak et al., 2002b) – it will be interesting to explore how non-auditory cues (Keleman et al., 2012; Zhang et al., 2016) modulate locomotor responses to song and whether multi-modal integration occurs in pC2 neurons or elsewhere. The absence of non-acoustic cues may explain the diversity of locomotor responses across strains in the FLyTRAP assay (Fig. S2). Using a naturalistic courtship assay, previous studies show that the same strains as the ones tested in FlyTRAP exhibit similar behaviors – males pattern their song in response to the female behavior and females change their locomotor speed to the natural courtship song similarly across all strains (Clemens et al., 2017; 2015; Coen et al., 2016; 2014).

In contrast to pulse song responses, the locomotor and singing responses for sine song in FLyTRAP were less sex-specific (Fig. 2E) and the behavioral tuning did not match well the conspecific song – very low frequencies never produced by males slowed NM91 females the most (Fig. 2A, B). This implies divergent roles for the two song modes and is consistent with previous studies (Eberl et al., 1997; F. V. Schilcher, 1976) – for instance sine song does not induce male-male courtship (Yoon et al., 2013). It has been suggested that pulse song may modulate sine song responses (F. V. Schilcher, 1976) but we did not detect strong serial interactions between the two song modes (Fig. S3G). Alternatively, responses to sine song may depend more strongly on the presence of male chemical cues (Billeter et al., 2009; Kurtovic et al., 2007) that are absent in the FLyTRAP assay. This is consistent with sine song being produced when the male is near the female (Coen et al., 2014) – that is, when these chemical cues are particularly strong.

### Pathways for detecting sine and pulse

Our behavioral and neuronal results suggest that pulse and sine song are processed in parallel pathways (Fig. 2E, 3C, F-H) but it is unclear as of yet how and where sounds are split into different streams. Sine and pulse can be separated based on spectral and temporal properties (Fig. S5). In fact, the frequency tuning in auditory receptor neurons (JON) and first-order auditory brain neurons (AMMC) may already be sufficient to separate the lower-frequency sine (150 Hz) from the higher-frequency pulse (>220 Hz) (Azevedo and Wilson, 2017; Ishikawa et al., 2017; Kamikouchi et al., 2009; Patella and Wilson, 2018; Yorozu et al., 2009). Temporal pattern could further discriminate pulse from sine by either suppressing responses to the sustained sine via adaptation or by tuning temporal integration such that the brief pulse stimuli fail to drive neuronal spiking. A complete mapping of auditory pathways and auditory activity throughout the *Drosophila* brain is required to identify where and how the neural selectivity for the different song modes arises.

Here, we have identified pC2 as one of the pathways driving responses to pulse song – pC2 tuning matches the behavioral tuning for pulse song (Fig. 3H, I), pC2 activation drives sex-specific responses to song (Fig. 5), and experience-dependent modulation of pC2 tuning matches the behavioral tuning (Fig. 6). Importantly, our data also indicate that pC2 neurons are not the only neurons used to detect pulse song, since the variability of pC2 neurons across stimuli and individuals does not account for the full behavioral variability (Fig. 3H, I, Fig. 4I-K). Interestingly, previous studies have implied pC1 as a pulse song detector (Zhou et al., 2015; 2014). Like pC2, pC1 exists in males and females (Rideout et al., 2010), and activation drives several courtship-related behaviors in males – including singing, male-male courtship, and aggression (Koganezawa et al., 2016; Kohatsu et al., 2011; Pan et al., 2012; Philipsborn et al., 2011; Zhou et al., 2015) – and also in females (Li et al., 2018; Rezával et al., 2016; Zhou et al., 2014). All previous studies have relied on imaging activity in the lateral junction (LJ) to show that pC1 preferentially responds to pulse song (Zhou et al., 2015; 2014). However, we show here that calcium responses of Dsx+ neurons in the LJ reflect the auditory activity of multiple Dsx+ cell types – and we detected auditory responses in the somas of pC2, pC1 (only in females) and pMN2 (a female only neuron) (Fig. 4). Because the number of auditory neurons within the pC2 cluster is much larger than for pC1 or pMN2 (Fig. 4C), and because tuning in pC2 somas matches the tuning in the LJ (Fig. 4E-H), we conclude that the LJ activity largely reflects pC2 responses. Nonetheless, we have not exhaustively assessed the match between the neuronal responses of female pC1 and pMN2 neurons and behavior. Those neurons may also be critical for the female’s response to pulse song, including behaviors not investigated here (such as oviposition (Kimura et al., 2015)).

### Inputs and outputs of pC2 neurons

pC2 neurons bind different properties of the pulse song to selectively signal the presence of conspecific pulse song – pC2 is tuned to several features of pulse song like pulse carrier frequency, pulse duration, and inter-pulse interval and a match in only one feature is not sufficient to strongly drive these neurons (Fig. 3, S4C-E). How this selectivity arises is as of yet unclear since systematic studies of tuning for multiple pulse song features in the early auditory pathway are missing. However, existing evidence suggests that pC2 may acquire its feature selectivity in a serial manner – via a cumulative sharpening of tuning for song features at successive stages of auditory processing (Kamikouchi et al., 2009; Yamada et al., 2018; Zhou et al., 2015; 2014). Auditory receptor neurons display diverse and specific band-pass tuning for carrier frequency (Ishikawa et al., 2017; Kamikouchi et al., 2009; Patella and Wilson, 2018; Yorozu et al., 2009) and first order auditory B1 neurons further sharpen frequency tuning via resonant conductances (Azevedo and Wilson, 2017). Likewise, peripheral responses are already weakly tuned for IPI (Clemens et al., 2018b; Ishikawa et al., 2017) and this tuning is further sharpened in downstream neurons (Vaughan et al., 2014; Zhou et al., 2015) through the interplay of excitation and inhibition (Yamada et al., 2018). This serial sharpening is similar to how selectivity for pulse song arises in crickets, in which a delay-line and coincidence detector mechanism produces broad selectivity for pulse duration and pulse pause which is subsequently sharpened in a downstream neuron (Schöneich et al., 2015). More direct readouts of the membrane voltage of auditory neurons in the fly brain are required to determine the biophysical mechanisms that generate song selectivity in pC2.

Similarly, the circuits downstream of pC2 neurons that control the diverse and sex-specific behaviors reported here remain to be identified. Our assessment of inter-individual variability in IPI preference revealed that most of the behavioral variability does not arise at the level of pC2 neurons (Fig. 4K). This suggests that variability in parallel or in downstream pathways strongly contributes to the locomotor tuning – pC2 activity is only one of multiple determinants of the behavior. This is consistent with optogenetic activation of pC2 being insufficient to produce the song-induced locomotor dynamics (Fig. S3, 5G,J). Still, our results demonstrate that pC2 drives sex-specific downstream circuits. pC2 neurons may connect directly with descending interneurons (DNs) (Cande et al., 2017; Namiki et al., 2017) that control motor behaviors. For example, pC2 activation in males drives pulse song production, followed by sine song production at stimulus offset (Fig. 5A). This behavior resembles that caused by pIP10 activation (Clemens et al., 2018a) – pIP10 is a male-only descending neuron (Philipsborn et al., 2011), but we don’t yet know if it directly connects with pC2 neurons. The fact that song responses are bi-directional – pulse song can induce both slowing and acceleration within each sex (Fig. 2A, B, S3) – implies that the sex-specificity of motor control is more than a simple re-routing from accelerating DNs in males to slowing DNs in females. Notably, song also promotes copulation, but we did not detect a significant effect of pC2 inactivation on copulation rates (Fig. S6O). This could be because our driver only labeled 1/3 of the auditory pC2 neurons or because pC2 activity does not inform the decision to mate. That is, parallel pathways may control song responses on different timescales: one pathway accumulates song information over timescales of minutes (Clemens et al., 2015; Ratcliff et al., 2016) and ultimately controls the mating decision; another, independent pathway controls behavioral responses to song on sub-second timescales, such as dynamic adjustments in locomotion and the production of courtship song.

### Modularity facilitates plasticity of behavioral responses to song

Our behavioral data suggest that some aspects of the sex-specificity of behavior arises after feature tuning. The pC2 neurons are selective for pulse song in both sexes (Fig. 3, 4) and drive locomotor responses with sex-specific dynamics or singing in males (Fig. 5). This is reminiscent of how sex-specific behaviors are driven to the male pheromone cVA in flies: shared detector neurons – olfactory receptor neurons and projection neurons in the antennal lobe – detect cVA in both sexes, and this information is then routed to sex-specific higher-order neurons in the lateral horn, which are thought to drive the different behaviors (Datta et al., 2008; Kohl et al., 2013; Ruta et al., 2010). This modular architecture with detectors of social signals being flexibly routed to different behavioral outputs is beneficial if these routes are plastic. For instance, here we show that social experience can shape male responses to song (similar to (Li et al., 2018)), along with tuning at the level of the neurons that detect the song (Fig. 6). During mating, males transfer a sex peptide to females (Yapici et al., 2008) that alters female behavioral responses to song from slowing to acceleration (Coen et al., 2014) – these effects may be mediated at the level of the motor circuits downstream of pC2, shifting pulse song responses in females to resemble those of males. Modularity also facilitates behavioral plasticity on evolutionary time scales since only one element – the feature detector – needs to change for behavioral tuning in both sexes to adapt to new songs that evolve during speciation (Capranica et al., 1973; Kostarakos et al., 2009). The identification of pC2 neurons as pulse song detectors is therefore likely to benefit future studies of the evolution of song recognition.

### pC2 neurons have a dual sensory and motor role

Unlike regular higher-order sensory neurons, which detect a sensory cue to drive different behaviors, pC2 neurons detect the cue whose production they drive (Fig. 3F, G, 5A-C). Such a dual sensory and motor role may guide social interactions and communication via imitation. In *Drosophila melanogaster*, hearing the song of other males induces a male to court and sing to other females and even males (Eberl et al., 1997; Yoon et al., 2013). This behavior may have originated because the song of another male indicates the presence of a female nearby.

Neurons with a dual sensory and motor roles are well-known from vertebrates (Mooney, 2014; Prather et al., 2008; Rizzolatti and Fogassi, 2014). For instance, “mirror” neurons are active during the production as well as the observation of a behavior and are thought to be crucial for imitation learning and communication between conspecifics (Rizzolatti and Arbib, 1998). Neurons with a sensorimotor correspondence in the brain of song birds are active during singing and hearing song, and these neurons are hypothesized to play a role in song learning (Mooney, 2014). Importantly, pC2 differs crucially from these instances in that it directly drives the production of the acoustic signal it detects (Fig. 5A-C). Because we recorded pC2 activity in passively listening males, we do not yet know whether pC2 is activated by sound in an actively singing male. If so, hearing its own song could induce self-stimulation and form a positive feedback loop to maintain courtship behavior by mediating persistent behavioral state-changes (Hoopfer et al., 2016). Alternatively, auditory inputs could be suppressed during singing via a corollary discharge (Poulet and Hedwig, 2003; Schneider et al., 2014), which would allow pC2 to maintain sensitivity to the song of other males to coordinate inter-male competition during singing. Additional studies of pC2 activity in behaving animals are required to fully understand how these pulse song detector neurons integrate into the acoustic communication behavior.

In summary, we show how the circuits that recognize song to drive diverse and sex-specific behavioral responses are organized in *Drosophila*: common detector neurons – pC2 – recognize pulse song in both males and females, and this identically processed information is then routed to drive multiple sex-specific behaviors. Similar principles may underlie the production of sex-specific behavioral responses to communication signals in other insects, song birds or mammals.

## Acknowledgements

Isabel D’Allesandro for help with playback behavioral experiments, Alex Hammons and Nofar Ozeri-Engelhard for help with dissections for immunostaining, Diego Pacheco for help with aligning the volumetric GCaMP scans, Robert Court and Doug Armstrong (http://www.virtualflybrain.org) for help with brain registration, David Stern, Ben Arthur and Barry Dickson for discussions during the development of the FLyTRAP assay, Kai Feng and Barry Dickson for sharing the design of their playback assay chamber, Bruce Baker, Stephen Goodwin, Gerry Rubin, Peter Andolfatto for gifts of flies, and Kristin Scott, Asif Ghazanfar, Tim Buschman, and members of the Murthy lab for feedback on the manuscript.

## Flies

The following fly lines were used in our study:

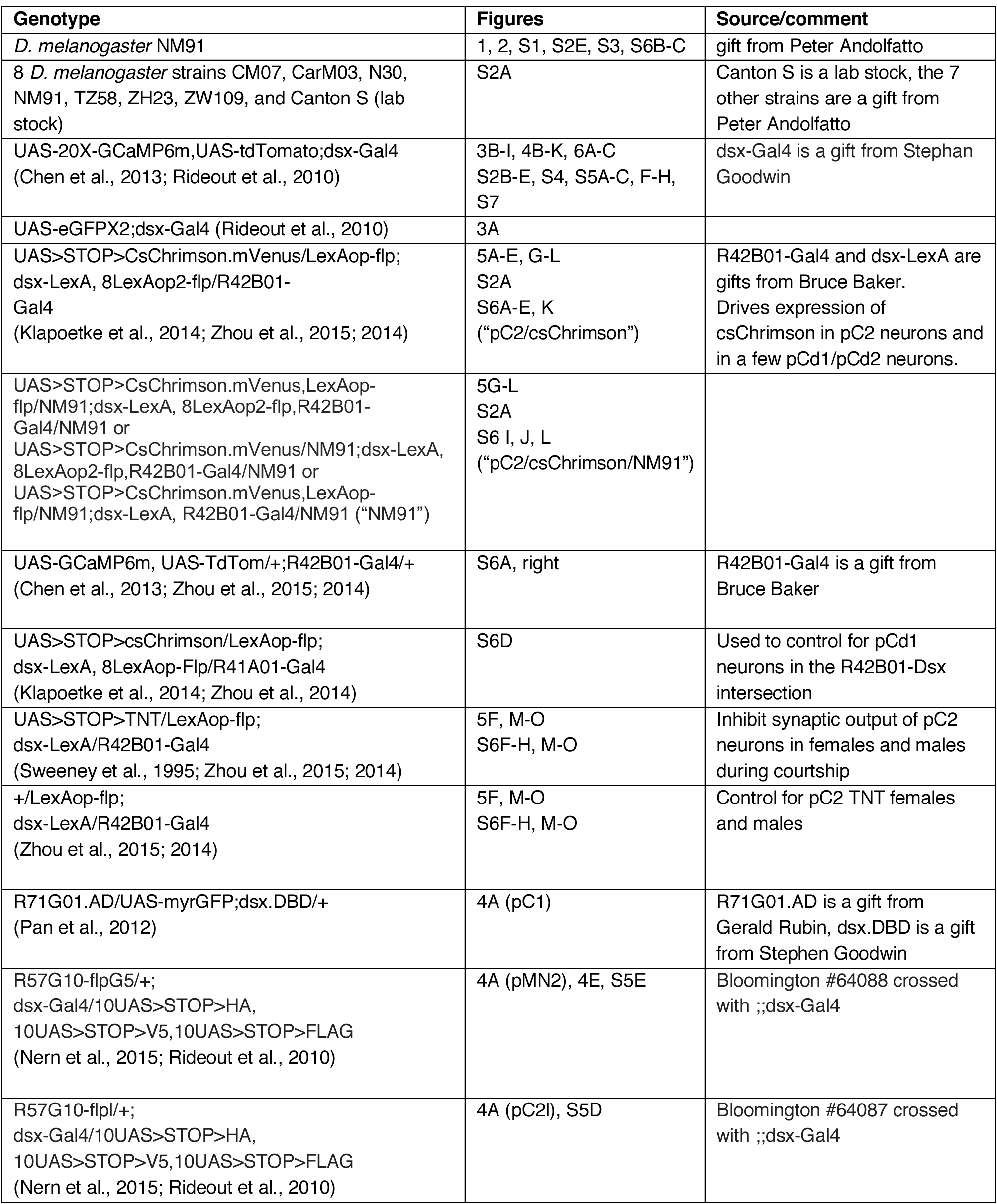

### FLyTRAP

Fly behavior was recorded with PointGrey cameras (FL3-U3-13Y3M-C or FL3-U3-13E4C-C). Grey color frames with a resolution of 1280×960 pixels were acquired at 30 frames per second using custom written software in python and saved as compressed videos. Sound representation was controlled using custom software written in Matlab. The sound stimuli were converted to an analog voltage signal using a National Instruments DAQ card (PCIe-6343). The signal was then amplified by a Samson s-amp headphone amp and used to drive a speaker (HiVi F6 6-1/2” Bass/Midrange). Sound intensity was calibrated as in (Clemens et al., 2015) by converting the voltage of a calibrated microphone (placed where the fly chambers would be during an experiment) to sound intensity and adjusting the sound amplification to match the target intensity. Sound and video where synchronized by placing into the camera’s field-of-view a 650nm LED whose brightness was controlled using a copy of the sound signal. The chamber consisted of an array of 12 small arenas (7 by 46 mm, made from red plastic) was placed in front of the loudspeaker (Movie S1). The arena floor consisted of plastic mesh to let sound into the chamber and the top was covered with a thin, translucent plastic sheet. Flies were illuminated using a white LED back light from below and a desk lamp from above.

### Playback experiments

Virgin male and female flies were isolated within 6 hours of eclosion and aged for 3-7 days prior to the experiments. Flies were raised at low density on a 12:12 dark:light cycle, at 25°C and 60% humidity. Flies were introduced gently into the chamber using an aspirator. Recordings were performed at 25°C and timed to start within 60 minutes of the incubator lights switching on to catch the morning activity peak. Stimulus playback was block-randomized to ensure that all stimuli within a set occur at the same overall rate throughout the stimulus. The stimulus set (e.g. five pulse trains with different IPIs, see Supplemental Table 1 for a list of all stimulus sets) was repeated for the duration of the experiment (2 hours). Stimuli were interleaved by 60 seconds of silence to reduce crosstalk between responses to subsequent stimulus presentations.

### Stimulus design

Sound was generated at a sampling frequency of 10 kHz using custom Matlab scripts. Sine song stimuli were created as pure tones of the specified frequency and intensity (typically 5mm/s). Pulse song was generated by arranging Gabor wavelets in trains interleaved by a specified pause. The Gabor wavelets were built by modulating the amplitude of a short sinusoidal using a Gaussian: *exp(-t^2^/(2σ^2^)) sin(2πf * t + ϕ),* where *f* is the pulse carrier frequency, *ϕ* is the phase of carrier, and *σ* is proportional to the pulse duration. The parameters for all stimuli used along with the behavioral responses obtained in FLyTRAP are listed in Supplemental Table 1.

### Analysis of FLyTRAP data

Fly positions where tracked using custom-written software. Briefly, the image background was estimated as the median of 500 frames spaced to cover the full video. Foreground pixels (corresponding to the fly body) were identified by thresholding the absolute values of the difference between each frame and the background estimate. The fly center position was then taken as the median of the position of all foreground pixels in each chamber. The sequence of fly positions across video frames was then converted into a time series using the light onset frames of the synchronization LED (indicating sound onset) as a reference. From the position time series fly speed was calculated and the speed traces where then aligned to stimulus onset for each trial. Base line speed was calculated as the average of the speed over an interval starting 30 seconds and ending 2 seconds before stimulus onset. Test speed was calculated over an interval starting at stimulus onset and ending 2 seconds after stimulus offset. Tuning curves were calculated as the difference between baseline speed and test speed for each trial, averaged over trials for each stimulus and animal. Speed traces were obtained by subtracting the baseline speed from the trace for each trial and averaging over trials for each stimulus and animal. All data (tuning curves, speed traces) are presented as mean +/-s.e.m. over flies. Code for stimulus generation, fly tracking and analysis of the locomotion data is available at https://github.com/murthylab/flytrapanalyses.

### Manual scoring of wing extension in FLyTRAP

To evaluate the number of flies that extend their wings upon playback of pulse or sine song, we manually scored wing extension in the videos using the VirtualDub software. For pulse song (see Movie S2), we scored 25 stimuli/fly, choosing trials randomly but ensuring that each IPI (16/36/56/76/96 ms) was scored 5 times/fly. To avoid bias, the scorer was blind to the IPI presented to the fly in each trial. A total of 120 male flies and 36 female flies were scored (3000 and 900 single-fly responses total for pulse song). We scored wing extension only when the wing was extended in the first 1/3 second following stimulus onset, and only when the wings where not extended during the 1 second before stimulus onset. For sine song (150Hz carrier frequency), 60 males were scored.

### Joint tuning for pulse duration and pulse pause

To visualize locomotor (Fig. S3E, F) and calcium (Fig. S4D,E) responses to pulse trains with different combinations of pulse duration and pulse pause we generated smooth surface plots using Matlab’s “scatteredInterpolant” function with the interpolation mode set to “natural”. The boundaries of the plots were set as follows: Pulse duration of zero corresponds to silence and the speed values were set to 0 since all speed traces are always base line subtracted. A pulse pause of zero corresponds to a continuous oscillation and we set the corresponding speed values to those obtained for a 4 second pure tone with a frequency of 250 Hz.

### Measurement of song features from natural song data

The inter-pulse interval (IPI) is given by the interval between the peaks of subsequent pulses in a pulse train. Pulse trains correspond to continuous sequences of pulses with IPIs smaller than 200ms. Measuring the pulse durations from natural song data is non-trivial since pulses vary in their shape and can be embedded in background noise. We quantified pulse duration by 1) calculating the envelope of each pulse using the Hilbert transform, 2) smoothing that envelope using a Gaussian window with a standard deviation of 2 ms, and 3) taking as the pulse duration the full width of the smoothed envelope at 20% of the maximum amplitude of the pulse. Pulse durations for artificial stimuli used in our pulse train were defined to be consistent with this method. Pulse carrier frequency is given by the center of mass of the amplitude spectrum of each pulse (Clemens et al., 2017). Sine carrier frequency was calculated as the peak frequency of the power spectrum of individual sine tones.

### PCA of speed traces

For the PCA of sex-specific responses to sound and optogenetic activation of pC2 (Fig. 5L) we collected male and female speed traces for all IPIs (Fig. 5D, F) and optogenetic activation levels (Fig. 5H, J) into a large matrix. Each speed trace was cut to include only the 10 seconds after sound onset and then normalized to have zero mean and unit variance.

### Optogenetic experiments

CsChrimson was expressed in pC2 neurons using an intersection between R42B01-Gal4 and dsx-LexA using two different genotypes (see table, pC2/csChrimson and pC2/csChrimson/NM91). 655nm light was emitted from a ring of 6 Tri-Star LEDs (LuxeonStar, SinkPAD-II 20mm Tri-Star Base) in FLyTRAP (Fig. 5D-K). Flies were fed with food that contained all-trans retinal for a minimum of three days post eclosion. Control flies were raised on regular fly food after eclosion. LED stimulation lasted four seconds with 60 seconds pause between stimuli, similar to the temporal pattern used for auditory stimulus delivery in FLyTRAP (1-5mW/cm^2^, 100Hz, duty cycle 0.5). Smaller intensities of 0.1-1wW/cm^2^ were not sufficient to drive changes in speed in the pC2/csChrimson/NM91 genotype (data not shown). To measure the amount of song driven by pC2 activation in solitary flies of the pC2/csChrimson and the pC2/csChrimson/NM91 genotype, we used a chamber whose floor was tiled with 16 microphones to allow recording of the song (Fig. 5A-C, Movie S7; see (Clemens et al., 2017)). The LED (627nm LEDs, LuxeonStar) was on for four seconds (frequency 25 Hz, duty cycle 0.5) and off for 60 seconds. For pC2/csChrimson, we tested three different light intensities (1.8, 9, and 13 mW/cm^2^) that were presented in 3 blocks of 18 trials. The order of the three blocks (light intensities) was randomized for each fly. pC2/csChrimson/NM91 was tested with 9mW/cm^2^ in 10 trials. Fly song was segmented as described previously (Arthur et al., 2013; Coen et al., 2014).

### pC2 inactivation in females and males during courtship

Tetanus neurotoxin light chain (TNT) (Sweeney et al., 1995) was used to block synaptic transmission in pC2 neurons in females and males. 3-7 days old virgin females or males (pC2-TNT: UAS>STOP>TNT/LexAop-flp; dsx-LexA/R42B01-Gal4, pC2-control: +/LexAop-flp; dsx-LexA/R42B01-Gal4) were paired with wild type flies (NM91) of the opposite sex, in a custom-built chamber designed to record fly song (∼25 mm diameter, tiled with 16 microphones; same setup as the one used for measuring optogenetic driven song). Flies were allowed to interact for 30 minutes, and the percent of flies copulated as a function of time was scored. A monochrome camera (Point Grey, FL3-U3-13Y3M) was used to record the fly behavior at 60 frames per second. Fly position was tracked offline and song was segmented as previously described (Arthur et al., 2013; Coen et al., 2014). We then calculated song statistics (e.g. amount of song or number of pulses per window) and female locomotion (average female speed) in windows of 60s with 30s overlap (Clemens et al., 2015). For the rank correlations between male song features and female speed (Fig. 5M-O), we binned the female speed values into 16 bins with the bin edges chosen such that each bin was populated by an equal amount of samples (see Fig. 5M) and calculated the rank correlation between the binned female speed and the average male song feature per bin. Changes in correlation between control and experimental flies (Fig. 5O) were analyzed using an ANCOVA model with independent slopes and intercepts. Significance was determined based on the p-value of the interaction term (model’s genotype by song-feature) after Bonferroni correction.

### Calcium imaging

Imaging experiments were performed on a custom built two-photon laser scanning microscope equipped with 5mm galvanometer mirrors (Cambridge Technology), an electro-optic modulator (M350-80LA-02 KD*P, Conoptics) to control the laser intensity, a piezoelectric focusing device (P-725, Physik Instrumente) for volumetric imaging, a Chameleon Ultra II Ti:Sapphire laser (Coherent) and a water immersion objective (Olympus XLPlan 25X, NA=1.05). The fluorescence signal collected by the objective was reflected by a dichroic mirror (FF685 Dio2, Semrock), filtered using a multiphoton short-pass emission filter (FF01-680/sp-25, Semrock), split by a dichroic mirror (FF555 Dio3, Semrock) into two channels, green (FF02-525/40-25, Semrock) and red (FF01-593/40-25, Semrock), and detected by GaAsP photo-multiplier tubes (H10770PA-40, Hamamatsu). Laser power (measured at the sample plane) was restricted to 15 mW. The microscope was controlled in Matlab using ScanImage 5.1 (Vidrio). Single plane calcium signals (Fig. 3C-I, 4F,G and pMN2 neuron in Fig 4C-E) were scanned at 8.5 Hz (256×256 pixels). Pixel size was ∼0.5μmX0.5μm when imaging the lateral junction or pC2l process and ∼0.25μmX0.25μm when imaging cell bodies in a single plan (Fig 4G and pMN2 in Fig. 4B-D). For volumetric scanning of cell bodies (Figs 4B-D, S5A), volumes were acquired at 0.5Hz (256*216, 20 planes, voxel size ∼ 0.34μm X 0.4μm X 1.5μm), scanning one group of cells at a time (pC1, pC2, pCd).

After surgery (opening of the head capsule to reveal the brain), flies were placed beneath the objective and perfusion saline was continuously delivered directly to the meniscus. Sound playback was controlled using custom written Matlab software (Clemens et al., 2018). The software also stopped and started the calcium imaging via a TTL pulse sent to ScanImage (“external hardware trigger” mode), and single frames were synchronized with stimulus by sensing a copy of the Y-galvo mirror to a National Instruments DAQ card (PCIe-6343) that controled the stimulus. The sound stimulus was generated at a sampling rate of 10kHz and sent by the DAQ card through an amplifier (Crown, D-75A) to a set of head phones (Koss, ‘The Plug’). A single ear plug was connected to one side of a plastic tube (outer-inner diameters 1/8’’-1/16’’) and the outer tube tip was positioned 2 mm away from the fly arista. Sound intensity was calibrated by measuring the sound intensity 2 mm away from the tube tip with a pre-calibrated microphone at a range of frequencies (100Hz-800Hz) and the output signal was corrected according to the measured intensities. The pause between stimulus representation was 25 seconds. A stimulus set (26-36 stimuli) was presented to each fly in a block-randomized order as in the playback experiments. Three blocks were presented for each fly. If the response decayed in the middle of a block (possibly because of drift in the z-axis), the whole block was discarded from the analysis. Typically, two full repetitions per fly were used for analysis.

Regions of interest (ROIs) for calcium response measurements (in the LJ, pC2 process and in single Dsx+ somata) were selected manually based on a z-projection of the tdTomato channel. ΔF/F of the GCaMP signal was calculated as (F(t)-F_0_)/ F_0_, where F_0_ is the mean fluorescence in the ROI in the 10 seconds preceding stimulus onset. Integral ΔF/F (Fig. 3D, F-I) and peak ΔF/F (Fig. 3F, inset) values were calculated in a window starting at sound stimulus onset and ending 25 seconds after sound stimulus offset. To compensate for differences in overall responsiveness across flies, we normalized ΔF/F values of each fly by dividing the integral or peak ΔF/F by the maximal value (of integral or peak ΔF/F) across all stimuli for that fly. For volumetric scanning (Fig. 4C-D, S5A) pulse song (250Hz, 16 pulse duration, 20 pulse pause), sine song (250 Hz) and broadband noise (100-900Hz) were presented 6 times each (in the order pulse-sine-noise, 6 blocks, duration of each stimulus 10 seconds with 20 of silence in between) for each group of neurons (pC1 or pC2). A cell was considered responsive to a given stimulus (pulse, sine or broadband noise) if the mean ΔF during the stimulus was higher than the mean ΔF in the 10 seconds before stimulus onset in 5/6 blocks. Each time series was first motion corrected using the rigid motion correction algorithm NoRMCorre (Pnevmatikakis and Giovannucci, 2017) taking the tdTomato signal as the reference image. Then, single cell bodies were drawn manually, by marking cell boundaries stack by stack. In some cases, mostly with male pC1 neurons, cell bodies were very packed, such that some ROIs we marked manually possibly included more than a single cell. The number of single cells reported from Ca imaging is therefore slightly underestimated.

### Light microscopy

Flies expressing GFP in Dsx+ neurons (UAS-eGFP2X; dsx-Gal4; Fig. 3A) and flies expressing CsChrimson.mVenus in pC2 neurons (R42B01-Gal4 intersected with dsx-LexA; Fig. S6) were immunostained and scanned in a confocal microscope. 2-4 day old flies were cold-anesthetized on ice, dissected in cold S2 insect medium (Sigma Aldrich, #S0146) and fixed for 30-40 minutes on a rotator at room temperature in 4% PFA in 0.3% PBTS (0.3% Triton in PBSX1), followed by 4×15 minutes washes in 0.3% PBTS and 30 minutes in blocking solution (5% normal goat serum in 0.3%PBTS). Brains were incubated over two nights at 4°C with primary antibody, washed with 0.3%PBT and incubated for two more nights at 4°C in secondary antibody, followed by washing (4×15 minutes in 0.3%PBTS and 4×20 minutes in PBS), and mounting with Vestashield for 2-7 days before imaging. Antibodies were diluted in blocking solution at the following concentrations: rabbit anti-GFP (Invitrogen #1828014; used against GFP and mVenus) 1:1000, mouse anti-Bruchpilot (nc82, DSHB AB2314866) 1:20, goat anti-rabbit Alexa Flour 488 (Invitrogen #1853312) 1:200, goat anti-mouse Alexa Flour 633 (Invitrogen #1906490) 1:200.

Stochastic labeling of Dsx+ neurons in the female brain (Fig 4A, E) was done using multi-color-flip-out (MCFO, (Nern et al., 2015)) with three different epitope tags (HA,V5,FLAG). We followed the JFRC FlyLight Protocol ‘IHC-MCFO’ (https://www.janelia.org/project-team/flylight/protocols) for the preparation of brains. Flp was induced using R5710C10 promotor-coding sequence fusions of the flpG5 and flpl. Flies were 4-7 days old when dissected. Flies were stored at 25°C. Confocal stacks were acquired with a white light laser confocal microscope (Leica TCS SP8 X) and a Leica objective (HC PL APO 20x/0.75 CS2). A high-resolution scan of a pC2 cell (Fig 4E) was performed with an oil immersion Leica objective (HC PL APO 63x/1.40 Oil CS2, fig 4E). Images were registered to the Janelia brain template (JFRC2) (Jenett et al., 2012) using vfbaligner (http://vfbaligner.inf.ed.ac.uk), which internally uses CMTK for registration (Rohlfing and Maurer, 2003). The images of the fly brain in Figs 4A and S5D were deposited by G. Jefferis (Jefferis, 2014). Image processing was performed in FIJI (Schindelin et al., 2012)

## References

Amézquita, A., Flechas, S.V., Lima, A.P., Gasser, H., Hödl, W., 2011. Acoustic interference and recognition space within a complex assemblage of dendrobatid frogs. Proc Natl Acad Sci U S A 108, 17058–17063. doi:10.1073/pnas.1104773108

Aranha, M.M., Herrmann, D., Cachitas, H., Neto-Silva, R.M., Dias, S., Vasconcelos, M.L., 2017. apterous Brain Neurons Control Receptivity to Male Courtship in Drosophila Melanogaster Females. Sci Rep 7, 46242. doi:10.1038/srep46242

Arthur, B.J., Sunayama-Morita, T., Coen, P., Murthy, M., Stern, D.L., 2013. Multi-channel acoustic recording and automated analysis of Drosophila courtship songs. BMC Biol 11, 11. doi:10.1186/1741-7007-11-11

Azevedo, A.W., Wilson, R.I., 2017. Active Mechanisms of Vibration Encoding and Frequency Filtering in Central Mechanosensory Neurons. Neuron 1–25. doi:10.1016/j.neuron.2017.09.004

Bennet-Clark, H.C., Ewing, A.W., 1969. Pulse interval as a critical parameter in the courtship song of Drosophila melanogaster. Animal Behaviour 17, 755–759. doi:10.1016/S0003-3472(69)80023-0

Bennet-Clark, H.C., Ewing, A.W., 1967. Stimuli provided by Courtship of Male Drosophila melanogaster. Nature 215, 669–671. doi:10.1038/215669a0

Billeter, J.-C., Atallah, J., Krupp, J.J., Millar, J.G., Levine, J.D., 2009. Specialized cells tag sexual and species identity in Drosophila melanogaster. Nature 461, 987–991. doi:10.1038/nature08495

Billeter, J.-C., Levine, J.D., 2013. Who is he and what is he to you? Recognition in Drosophila melanogaster. Current Opinion in Neurobiology 23, 17–23. doi:10.1016/j.conb.2012.08.009

Bizley, J.K., Cohen, Y.E., 2013. The what, where and how of auditory-object perception. Nature Reviews Neuroscience 14, 693–707. doi:10.1038/nrn3565

Blankers, T., Hennig, R.M., Gray, D.A., 2015. Conservation of multivariate female preference functions and preference mechanisms in three species of trilling field crickets. Journal of Evolutionary Biology n/a–n/a. doi:10.1111/jeb.12599

Bussell, J.J., Yapici, N., Zhang, S.X., Dickson, B.J., Vosshall, L.B., 2014. Abdominal-B Neurons Control Drosophila Virgin Female Receptivity. Current Biology 24, 1584–1595. doi:10.1016/j.cub.2014.06.011

Cachero, S., Ostrovsky, A.D., Yu, J.Y., Dickson, B.J., Jefferis, G.S.X.E., 2010. Sexual dimorphism in the fly brain. Current biology : CB 20, 1589–1601. doi:10.1016/j.cub.2010.07.045

Cande, J., Berman, G.J., Namiki, S., Qiu, J., Korff, W., Card, G., Shaevitz, J.W., Stern, D.L., 2017. Optogenetic dissection of descending behavioral control in Drosophila. bioRxiv 1– 50. doi:10.1101/230128

Capranica, R.R., Frishkopf, L.S., Nevo, E., 1973. Encoding of Geographic Dialects in the Auditory System of the Cricket Frog. Science 182, 1272–1275. doi:10.1126/science.182.4118.1272

Chen, T.-W., Wardill, T.J., Sun, Y., Pulver, S.R., Renninger, S.L., Baohan, A., Schreiter, E.R., Kerr, R.A., Orger, M.B., Jayaraman, V., Looger, L.L., Svoboda, K., Kim, D.S., 2013. Ultrasensitive fluorescent proteins for imaging neuronal activity. Nature 499, 295– 300. doi:10.1038/nature12354

Clemens, J., Coen, P., Roemschied, F., Pereira, T., Mazumder, D., Pacheco, D., Murthy, M., 2017. Discovery of a new song mode in Drosophila reveals hidden structure in the sensory and neural drivers of behavior. bioRxiv 221044. doi:10.1101/221044

Clemens, J., Coen, P., Roemschied, F.A., Pereira, T.D., Mazumder, D., Aldarondo, D.E., Pacheco, D.A., Murthy, M., 2018a. Discovery of a New Song Mode in Drosophila Reveals Hidden Structure in the Sensory and Neural Drivers of Behavior. Current Biology 28, 2400–2412.e6. doi:10.1016/j.cub.2018.06.011

Clemens, J., Girardin, C.C., Coen, P., Guan, X.-J., Dickson, B.J., Murthy, M., 2015. Connecting Neural Codes with Behavior in the Auditory System of Drosophila. Neuron 87, 1332–1343. doi:10.1016/j.neuron.2015.08.014

Clemens, J., Hennig, R.M., 2013. Computational principles underlying the recognition of acoustic signals in insects. Journal of Computational Neuroscience 35, 75–85. doi:10.1007/s10827-013-0441-0

Clemens, J., Ozeri-Engelhard, N., Murthy, M., 2018b. Fast intensity adaptation enhances the encoding of sound in Drosophila. Nat Commun 9, 134. doi:10.1038/s41467-017-02453-9

Clyne, J.D., Miesenböck, G., 2008. Sex-specific control and tuning of the pattern generator for courtship song in Drosophila. Cell 133, 354–363. doi:10.1016/j.cell.2008.01.050

Coen, P., Clemens, J., Weinstein, A.J., Pacheco, D.A., Deng, Y., Murthy, M., 2014. Dynamic sensory cues shape song structure in Drosophila. Nature 507, 233–237. doi:10.1038/nature13131

Coen, P., Xie, M., Clemens, J., Murthy, M., 2016. Sensorimotor Transformations Underlying Variability in Song Intensity during Drosophila Courtship. Neuron 89, 629–644. doi:10.1016/j.neuron.2015.12.035

Cook, R.M., 1973. Courtship processing in Drosophila melanogaster. II. An adaptation to selection for receptivity to wingless males. Animal Behaviour 21, 349–358. doi:10.1016/S0003-3472(73)80077-6

Crossley, S.A., Bennet-Clark, H.C., Evert, H.T., 1995. Courtship song components affect male and female Drosophila differently. Animal Behaviour 50, 827–839. doi:10.1016/0003-3472(95)80142-1

Datta, S.R., Vasconcelos, M.L., Ruta, V., Luo, S., Wong, A., Demir, E., Flores, J., Balonze, K., Dickson, B.J., Axel, R., 2008. The Drosophila pheromone cVA activates a sexually dimorphic neural circuit. Nature 452, 473–477. doi:10.1038/nature06808

Dicarlo, J.J., Zoccolan, D., Rust, N.C., 2012. How Does the Brain Solve Visual Object Recognition? Neuron 73, 415–434. doi:10.1016/j.neuron.2012.01.010

Dulac, C., Wagner, S., 2006. Genetic Analysis of Brain Circuits Underlying Pheromone Signaling. Annu. Rev. Genet. 40, 449–467. doi:10.1146/annurev.genet.39.073003.093937

Eberl, D.F., Duyk, G.M., Perrimon, N., 1997. A genetic screen for mutations that disrupt an auditory response in Drosophila melanogaster. Proceedings of the National Academy of Sciences of the United States of America 94, 14837–14842. doi:10.1146/annurev.neuro.20.1.567

Ellis, L.B., Kessler, S., 1975. Differential posteclosion housing experiences and reproduction in Drosophila. Animal Behaviour 23, 949–952. doi:10.1016/0003-3472(75)90119-0

Fan, P., Manoli, D.S., Ahmed, O.M., Chen, Y., Agarwal, N., Kwong, S., Cai, A.G., Neitz, J., Renslo, A., Baker, B.S., Shah, N.M., 2013. Genetic and Neural Mechanisms that Inhibit Drosophila from Mating with Other Species. Cell 154, 89–102. doi:10.1016/j.cell.2013.06.008

Fortune, E.S., Rodríguez, C., Li, D., Ball, G.F., Coleman, M.J., 2011. Neural mechanisms for the coordination of duet singing in wrens. Science 334, 666–670. doi:10.1126/science.1209867

Gentner, T.Q., 2008. Temporal scales of auditory objects underlying birdsong vocal recognition. The Journal of the Acoustical Society of America 124, 1350–1359. doi:10.1121/1.2945705

Gentner, T.Q., Margoliash, D., 2003. Neuronal populations and single cells representing learned auditory objects. Nature 424, 669–674. doi:10.1038/nature01731

Gerhardt, C.H., Huber, F., 2002. Acoustic Communication in Insects and Anurans. University Of Chicago Press.

Griffiths, T.D., Warren, J.D., 2004. What is an auditory object? Nature Reviews Neuroscience 5, 887–892. doi:10.1038/nrn1538

Haga, S., Hattori, T., Sato, T., Sato, K., Matsuda, S., Kobayakawa, R., Sakano, H., Yoshihara, Y., Kikusui, T., Touhara, K., 2010. The male mouse pheromone ESP1 enhances female sexual receptive behaviour through a specific vomeronasal receptor. Nature 466, 118–122. doi:10.1038/nature09142

Helversen, von, D., Helversen, von, O., 1997. Recognition of sex in the acoustic communication of the grasshopper Chorthippus biguttulus (Orthoptera, Acrididae). Journal of Comparative Physiology A: Neuroethology, Sensory, Neural, and Behavioral Physiology 180, 373–386. doi:10.1007/s003590050056

Hennig, R.M., Blankers, T., Gray, D.A., 2016. Divergence in male cricket song and female preference functions in three allopatric sister species. Journal of Comparative Physiology A: Neuroethology, Sensory, Neural, and Behavioral Physiology 202, 347– 360. doi:10.1007/s00359-016-1083-2

Hennig, R.M., Heller, K.-G., Clemens, J., 2014. Time and timing in the acoustic recognition system of crickets. Frontiers in Physiology 5. doi:10.3389/fphys.2014.00286

Hoopfer, E.D., Jung, Y., Inagaki, H.K., Rubin, G.M., Anderson, D.J., Ramaswami, M., 2016. P1 interneurons promote a persistent internal state that enhances inter-male aggression in Drosophila. eLife 4, e11346. doi:10.7554/eLife.11346

Ishii, K.K., Osakada, T., Mori, H., Miyasaka, N., Yoshihara, Y., Miyamichi, K., Touhara, K., 2017. A Labeled-Line Neural Circuit for Pheromone-Mediated Sexual Behaviors in Mice. Neuron 95, 123–137.e8. doi:10.1016/j.neuron.2017.05.038

Ishikawa, Y., Okamoto, N., Nakamura, M., Kim, H., Kamikouchi, A., 2017. Anatomic and Physiologic Heterogeneity of Subgroup-A Auditory Sensory Neurons in Fruit Flies. Front. Neural Circuits 11, 46. doi:10.3389/fncir.2017.00046

Ito, K., Shinomiya, K., Ito, M., Armstrong, J.D., Boyan, G., Hartenstein, V., Harzsch, S., Heisenberg, M., Homberg, U., Jenett, A., Keshishian, H., Restifo, L.L., Rössler, W., Simpson, J.H., Strausfeld, N.J., Strauss, R., Vosshall, L.B., 2014. A Systematic Nomenclature for the Insect Brain. Neuron 81, 755–765. doi:10.1016/j.neuron.2013.12.017

Jazayeri, M., Afraz, A., 2017. Navigating the Neural Space in Search of the Neural Code. Neuron 93, 1003–1014. doi:10.1016/j.neuron.2017.02.019

Kamikouchi, A., Inagaki, H.K., Effertz, T., Hendrich, O., Fiala, A., Göpfert, M.C., Ito, K., 2009. The neural basis of Drosophila gravity-sensing and hearing. Nature 458, 165–171. doi:10.1038/nature07810

Keleman, K., Vrontou, E., Krüttner, S., Yu, J.Y., Kurtovic-Kozaric, A., Dickson, B.J., 2012. Dopamine neurons modulate pheromone responses in Drosophila courtship learning. Nature. doi:10.1038/nature11345

Kelley, D.B., 2003. Sexually Dimorphic Behaviors. http://dx.doi.org/10.1146/annurev.ne.11.030188.001301 11, 225–251.

Kimura, K.-I., Sato, C., Koganezawa, M., Yamamoto, D., 2015. Drosophila Ovipositor Extension in Mating Behavior and Egg Deposition Involves Distinct Sets of Brain Interneurons. PLoS ONE 10, e0126445. doi:10.1371/journal.pone.0126445

Klapoetke, N.C., Murata, Y., Kim, S.S., Pulver, S.R., Birdsey-Benson, A., Cho, Y.K., Morimoto, T.K., Chuong, A.S., Carpenter, E.J., Tian, Z., Wang, J., Xie, Y., Yan, Z., Zhang, Y., Chow, B.Y., Surek, B., Melkonian, M., Jayaraman, V., Constantine-Paton, M., Wong, G.K.-S., Boyden, E.S., 2014. Independent optical excitation of distinct neural populations. Nat. Methods 11, 338–346. doi:10.1038/nmeth.2836

Koganezawa, M., Kimura, K.-I., Yamamoto, D., 2016. The Neural Circuitry that Functions as a Switch for Courtship versus Aggression in Drosophila Males. Current Biology 26, 1395–1403. doi:10.1016/j.cub.2016.04.017

Kohatsu, S., Koganezawa, M., Yamamoto, D., 2011. Female contact activates male-specific interneurons that trigger stereotypic courtship behavior in Drosophila. Neuron 69, 498– 508. doi:10.1016/j.neuron.2010.12.017

Kohatsu, S., Yamamoto, D., 2015. Visually induced initiation of Drosophila innate courtship-like following pursuit is mediated by central excitatory state. Nat Commun 6, 6457. doi:10.1038/ncomms7457

Kohl, J., Ostrovsky, A.D., Frechter, S., Jefferis, G.S.X.E., 2013. A Bidirectional Circuit Switch Reroutes Pheromone Signals in Male and Female Brains. Cell 155, 1610–1623. doi:10.1016/j.cell.2013.11.025

Konishi, M., 1985. Birdsong: From Behavior to Neuron. Annu. Rev. Neurosci. 8, 125–170. doi:10.1146/annurev.neuro.8.1.125

Kostarakos, K., Hennig, M.R., Römer, H., 2009. Two matched filters and the evolution of mating signals in four species of cricket. Frontiers in Zoology 6, 22. doi:10.1186/1742-9994-6-22

Kurtovic, A., Widmer, A., Dickson, B.J., 2007. A single class of olfactory neurons mediates behavioural responses to a Drosophila sex pheromone. Nature 446, 542–546. doi:10.1038/nature05672

Lehnert, B.P., Baker, A.E., Gaudry, Q., Chiang, A.-S., Wilson, R.I., 2013. Distinct Roles of TRP Channels in Auditory Transduction and Amplification in Drosophila. Neuron 77, 115–128. doi:10.1016/j.neuron.2012.11.030

Li, X., Ishimoto, H., Kamikouchi, A., 2018. Auditory experience controls the maturation of song discrimination and sexual response in Drosophila. eLife 7, e34348. doi:10.7554/eLife.34348

Marlin, B.J., Mitre, M., D’Amour, J.A., Chao, M.V., Froemke, R.C., 2015. Oxytocin enables maternal behaviour by balancing cortical inhibition. Nature 520, 499–504. doi:10.1038/nature14402

Mooney, R., 2014. Auditory–vocal mirroring in songbirds. Philosophical Transactions of the Royal Society B: Biological Sciences 369, 20130179–399. doi:10.1098/rstb.2013.0179

Nagel, K.I., Wilson, R.I., 2011. Biophysical mechanisms underlying olfactory receptor neuron dynamics. Nature neuroscience 14, 208–216. doi:10.1038/nn.2725

Namiki, S., Dickinson, M.H., Wong, A.M., Korff, W., Card, G.M., 2017. The functional organization of descending sensory-motor pathways in Drosophila. bioRxiv 231696. doi:10.1101/231696

Nern, A., Pfeiffer, B.D., Rubin, G.M., 2015. Optimized tools for multicolor stochastic labeling reveal diverse stereotyped cell arrangements in the fly visual system. Proc Natl Acad Sci U S A 112, E2967–76. doi:10.1073/pnas.1506763112

Pan, Y., Meissner, G.W., Baker, B.S., 2012. Joint control of Drosophila male courtship behavior by motion cues and activation of male-specific P1 neurons. Proc Natl Acad Sci U S A 109, 10065–10070. doi:10.1073/pnas.1207107109

Patella, P., Wilson, R.I., 2018. Functional Maps of Mechanosensory Features in the Drosophila Brain. Current Biology 0. doi:10.1016/j.cub.2018.02.074

Philipsborn, von, A.C., Liu, T., Yu, J.Y., Masser, C., Bidaye, S.S., Dickson, B.J., 2011. Neuronal control of Drosophila courtship song. Neuron 69, 509–522. doi:10.1016/j.neuron.2011.01.011

Poulet, J.F.A., Hedwig, B., 2003. Corollary discharge inhibition of ascending auditory neurons in the stridulating cricket. The Journal of neuroscience : the official journal of the Society for Neuroscience 23, 4717–4725.

Prather, J.F., Peters, S., Nowicki, S., Mooney, R., 2008. Precise auditory–vocal mirroring in neurons for learned vocal communication. Nature 451, 305–310. doi:10.1038/nature06492

Ratcliff, R., Smith, P.L., Brown, S.D., McKoon, G., 2016. Diffusion Decision Model: Current Issues and History. Trends Cogn. Sci. (Regul. Ed.) 20, 260–281. doi:10.1016/j.tics.2016.01.007

Remedios, R., Kennedy, A., Zelikowsky, M., Grewe, B.F., Schnitzer, M.J., Anderson, D.J., 2017. Social behaviour shapes hypothalamic neural ensemble representations of conspecific sex. Nature 550, 388–392. doi:10.1038/nature23885

Rezával, C., Pattnaik, S., Pavlou, H.J., Nojima, T., Brüggemeier, B., D’Souza, L.A.D., Dweck, H.K.M., Goodwin, S.F., 2016. Activation of Latent Courtship Circuitry in the Brain of Drosophila Females Induces Male-like Behaviors. Current Biology 26, 2508– 2515. doi:10.1016/j.cub.2016.07.021

Riabinina, O., Dai, M., Duke, T., Albert, J.T., 2011. Active process mediates species-specific tuning of Drosophila ears. Current biology : CB 21, 658–664. doi:10.1016/j.cub.2011.03.001

Rideout, E.J., Dornan, A.J., Neville, M.C., Eadie, S., Goodwin, S.F., 2010. Control of sexual differentiation and behavior by the doublesex gene in Drosophila melanogaster. Nature neuroscience 13, 458–466. doi:10.1038/nn.2515

Rizzolatti, G., Arbib, M.A., 1998. Language within our grasp. Trends in Neurosciences 21, 188–194. doi:10.1016/S0166-2236(98)01260-0

Rizzolatti, G., Fogassi, L., 2014. The mirror mechanism: recent findings and perspectives. Philosophical Transactions of the Royal Society B: Biological Sciences 369, 20130420– 20130420. doi:10.1098/rstb.2013.0420

Robinett, C.C., Vaughan, A.G., Knapp, J.M., Baker, B.S., 2010. Sex and the single cell. II. There is a time and place for sex. PLoS Biology 8, e1000365. doi:10.1371/journal.pbio.1000365.g009

Ronacher, B., Ronacher, B., Hennig, R.M., Clemens, J., 2014. Computational principles underlying recognition of acoustic signals in grasshoppers and crickets. J Comp Physiol A 201, 61–71. doi:10.1007/s00359-014-0946-7

Rosenthal, G.G., Ryan, M.J., 2011. Conflicting preferences within females: sexual selection versus species recognition. Biol. Lett. 7, 525–527. doi:10.1098/rsbl.2011.0027

Ruta, V., Datta, S.R., Vasconcelos, M.L., Freeland, J., Looger, L.L., Axel, R., 2010. A dimorphic pheromone circuit in Drosophila from sensory input to descending output. Nature 468, 686–690. doi:10.1038/nature09554

Ryan, M.J., Cummings, M.E., 2013. Perceptual Biases and Mate Choice. Annu. Rev. Ecol. Evol. Syst. 44, 437–459. doi:10.1146/annurev-ecolsys-110512-135901

Ryan, M.J., Phelps, S.M., Rand, A.S., 2001. How evolutionary history shapes recognition mechanisms. Trends Cogn. Sci. (Regul. Ed.) 5, 143–148.

Rybak, F., Aubin, T., Moulin, B., Jallon, J.-M., 2002a. Acoustic communication in Drosophila melanogaster courtship: Are pulse-and sine-song frequencies important for courtship success? Canadian journal of zoology 80, 987–996.

Rybak, F., Sureau, G., Aubin, T., 2002b. Functional coupling of acoustic and chemical signals in the courtship behaviour of the male Drosophila melanogaster. Proc. R. Soc. B 269, 695–701. doi:10.1098/rspb.2001.1919

Schilcher, F.V., 1976. The function of pulse song and sine song in the courtship of Drosophila melanogaster. Animal Behaviour 24, 622–625.

Schilcher, Von, F., 1976. The role of auditory stimuli in the courtship of Drosophila melanogaster. Animal Behaviour 24, 18–26. doi:10.1016/S0003-3472(76)80095-4

Schneider, D.M., Nelson, A., Mooney, R., 2014. A synaptic and circuit basis for corollary discharge in the auditory cortex. Nature. doi:10.1038/nature13724

Schöneich, S., Kostarakos, K., Hedwig, B., 2015. An auditory feature detection circuit for sound pattern recognition. Science Advances 1, e1500325–e1500325. doi:10.1126/sciadv.1500325

Shirangi, T.R., Stern, D.L., Truman, J.W., 2013. Motor control of Drosophila courtship song. Cell Reports 5, 678–686. doi:10.1016/j.celrep.2013.09.039

Stowers, L., Logan, D.W., 2010. Sexual dimorphism in olfactory signaling. Current Opinion in Neurobiology 20, 770–775. doi:10.1016/j.conb.2010.08.015

Sweeney, S.T., Broadie, K., Keane, J., Niemann, H., O’Kane, C.J., 1995. Targeted expression of tetanus toxin light chain in Drosophila specifically eliminates synaptic transmission and causes behavioral defects. Neuron 14, 341–351. doi:10.1016/0896-6273(95)90290-2

Talyn, B.C., Dowse, H.B., 2004. The role of courtship song in sexual selection and species recognition by female Drosophila melanogaster. Animal Behaviour 68, 1165–1180. doi:10.1016/j.anbehav.2003.11.023

Tinbergen, N., 1989. The Study of Instinct. Oxford University Press.

Tompkins, L., Gross, A.C., Hall, J.C., Gailey, D.A., Siegel, R.W., 1982. The role of female movement in the sexual behavior ofDrosophila melanogaster. Behav Genet 12, 295– 307. doi:10.1007/BF01067849

Tsao, D.Y., Livingstone, M.S., 2008. Mechanisms of Face Perception. Annu. Rev. Neurosci. 31, 411–437. doi:10.1146/annurev.neuro.30.051606.094238

Vaughan, A.G., Zhou, C., Manoli, D.S., Baker, B.S., 2014. Neural Pathways for the Detection and Discrimination of Conspecific Song in D. melanogaster. Current Biology. doi:10.1016/j.cub.2014.03.048

Versteven, M., Vanden Broeck, L., Geurten, B., Zwarts, L., Decraecker, L., Beelen, M., Göpfert, M.C., Heinrich, R., Callaerts, P., 2017. Hearing regulates Drosophila aggression. Proc Natl Acad Sci U S A 114, 1958–1963. doi:10.1073/pnas.1605946114

Wang, L., Anderson, D.J., 2010. Identification of an aggression-promoting pheromone and its receptor neurons in Drosophila. Nature 463, 227–231. doi:10.1038/nature08678

Yamada, D., Ishimoto, H., Li, X., Kohashi, T., Ishikawa, Y., Kamikouchi, A., 2018. GABAergic Local Interneurons Shape Female Fruit Fly Response to Mating Songs. J. Neurosci. 38, 4329–4347. doi:10.1523/JNEUROSCI.3644-17.2018

Yamamoto, D., Yamamoto, D., Koganezawa, M., Koganezawa, M., 2013. Genes and circuits of courtship behaviour in Drosophila males. Nature Reviews Neuroscience 14, 681–692. doi:10.1038/nrn3567

Yang, C.F., Shah, N.M., 2014. Representing Sex in the Brain, One Module at a Time. Neuron 82, 261–278. doi:10.1016/j.neuron.2014.03.029

Yapici, N., Kim, Y.J., Ribeiro, C., Dickson, B.J., 2008. A receptor that mediates the post-mating switch in Drosophila reproductive behaviour. Nature.

Yoon, J., Matsuo, E., Yamada, D., Mizuno, H., Morimoto, T., Miyakawa, H., Kinoshita, S., Ishimoto, H., Kamikouchi, A., 2013. Selectivity and Plasticity in a Sound-Evoked Male-Male Interaction in Drosophila. PLoS ONE 8, e74289. doi:10.1371/journal.pone.0074289

Yorozu, S., Wong, A., Fischer, B.J., Dankert, H., Kernan, M.J., Kamikouchi, A., Ito, K., Anderson, D.J., 2009. Distinct sensory representations of wind and near-field sound in the Drosophila brain. Nature 458, 201–205. doi:10.1038/nature07843

Yu, J.Y., Kanai, M.I., Demir, E., Jefferis, G.S.X.E., Dickson, B.J., 2010. Cellular organization of the neural circuit that drives Drosophila courtship behavior. Current biology : CB 20, 1602–1614. doi:10.1016/j.cub.2010.08.025

Zhang, S.X., Rogulja, D., Crickmore, M.A., 2016. Dopaminergic Circuitry Underlying Mating Drive. Neuron 91, 168–181. doi:10.1016/j.neuron.2016.05.020

Zhou, C., Franconville, R., Vaughan, A.G., Robinett, C.C., Jayaraman, V., Baker, B.S., 2015. Central neural circuitry mediating courtship song perception in male Drosophila. eLife 4, 11. doi:10.7554/eLife.08477

Zhou, C., Pan, Y., Robinett, C.C., Meissner, G.W., Baker, B.S., 2014. Central Brain Neurons Expressing doublesex Regulate Female Receptivity in Drosophila. Neuron 83, 149–163. doi:10.1016/j.neuron.2014.05.038

## References

Chen, T.-W., Wardill, T.J., Sun, Y., Pulver, S.R., Renninger, S.L., Baohan, A., Schreiter, E.R., Kerr, R.A., Orger, M.B., Jayaraman, V., Looger, L.L., Svoboda, K., Kim, D.S., 2013. Ultrasensitive fluorescent proteins for imaging neuronal activity. Nature 499, 295–300. doi:10.1038/nature12354

Clemens, J., Ozeri-Engelhard, N., Murthy, M., 2018. Fast intensity adaptation enhances the encoding of sound in Drosophila. Nat Commun 9, 134. doi:10.1038/s41467-017-02453-9

Jefferis, G.S.X.E., 2014. JFRC2 Template Brain. doi:10.5281/zenodo.10567

Jenett, A., Rubin, G.M., Ngo, T.-T.B., Shepherd, D., Murphy, C., Dionne, H., Pfeiffer, B.D., Cavallaro, A., Hall, D., Jeter, J., Iyer, N., Fetter, D., Hausenfluck, J.H., Peng, H., Trautman, E.T., Svirskas, R.R., Myers, E.W., Iwinski, Z.R., Aso, Y., DePasquale, G.M., Enos, A., Hulamm, P., Lam, S.C.B., Li, H.-H., Laverty, T.R., Long, F., Qu, L., Murphy, S.D., Rokicki, K., Safford, T., Shaw, K., Simpson, J.H., Sowell, A., Tae, S., Yu, Y., Zugates, C.T., 2012. A GAL4-Driver Line Resource for Drosophila Neurobiology. Cell Reports 2, 991–1001. doi:10.1016/j.celrep.2012.09.011

Pnevmatikakis, E.A., Giovannucci, A., 2017. NoRMCorre: An online algorithm for piecewise rigid motion correction of calcium imaging data. Journal of Neuroscience Methods 291, 83–94. doi:10.1016/j.jneumeth.2017.07.031

Rohlfing, T., Maurer, C.R., 2003. Nonrigid image registration in shared-memory multiprocessor environments with application to brains, breasts, and bees. IEEE Trans Inf Technol Biomed 7, 16–25.

Schindelin, J., Arganda-Carreras, I., Frise, E., Kaynig, V., Longair, M., Pietzsch, T., Preibisch, S., Rueden, C., Saalfeld, S., Schmid, B., Tinevez, J.-Y., White, D.J., Hartenstein, V., Eliceiri, K., Tomancak, P., Cardona, A., 2012. Fiji: an open-source platform for biological-image analysis. Nat. Methods 9, 676–682. doi:10.1038/nmeth.2019

