## Supplemental Data 1 for "Shared Song Detector Neurons in *Drosophila* Male and Female Brains Drive Sex-Specific Behaviors"

#### Supplemental Figures

**Figure S1**

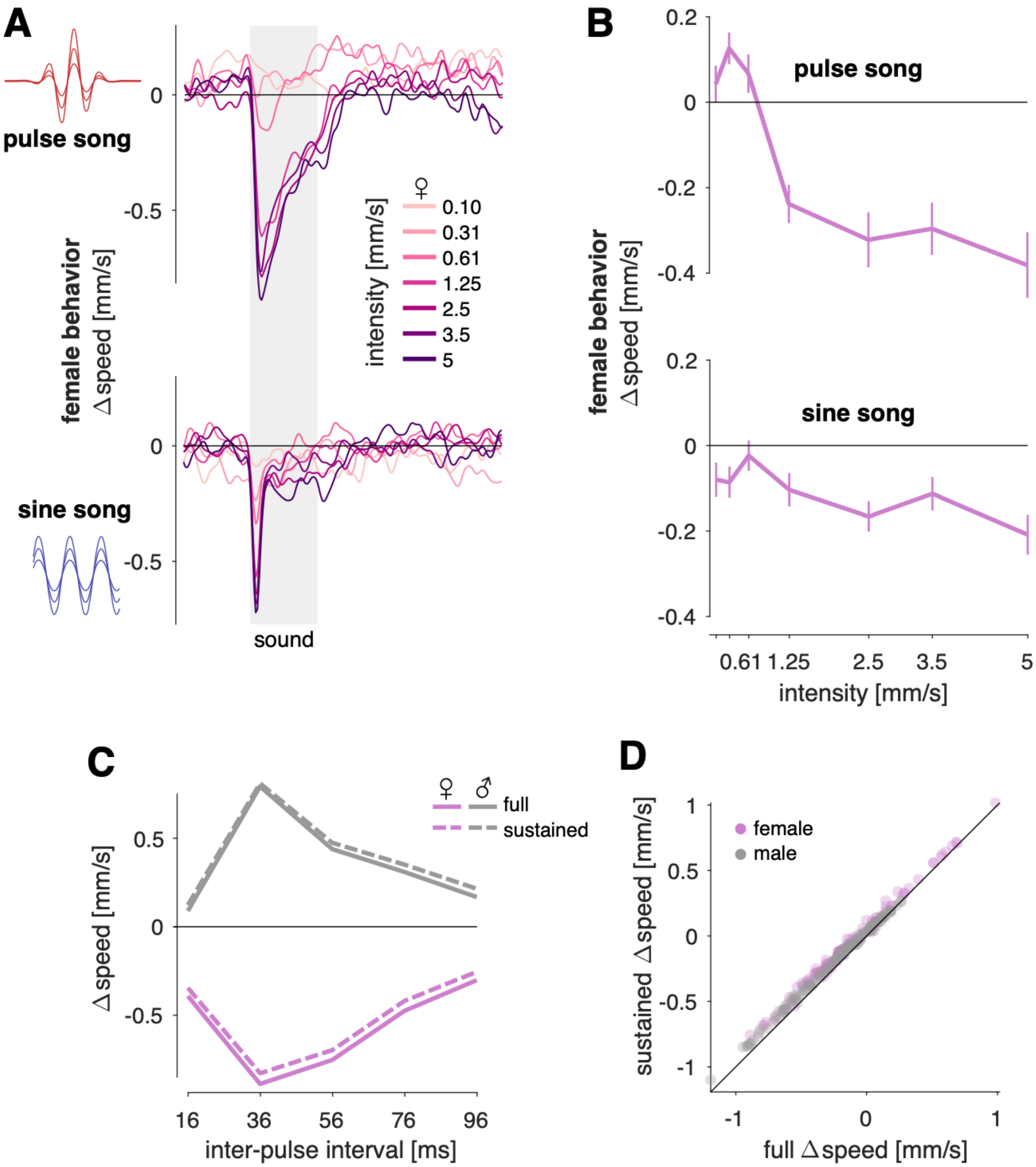

**Figure S1. Related to Figure 1.**
**A** Female locomotor responses for pulse trains (36ms IPI stimulus) and sine tones (100Hz tones) of different intensities (color coded, see legend). Intensity is given in mm/s since flies are sensitive to the particle velocity of sound, not sound pressure.
**B** Speed tuning curves for the traces in A obtained by averaging the speed in the six seconds following stimulus onset. For pulse song, responses are weak for intensities <1mm/s and don't change much beyond that. While sine responses are weak, there is a tendency for speed to change more for louder sine tones. Lines and error bars indicate mean±s.e.m. over ~120 flies.
**C** Excluding the transient response component only negligibly affects behavioral tuning curves. Shown are IPI tuning curves for males (gray) and females (magenta) obtained by averaging different epochs of the speed traces. The full response (solid lines) corresponds to the average, base-line subtracted speed in the 6 seconds following sound onset. For the sustained response (dashed lines), we start integration of the speed traces not at sound onset but 500ms into the sound. Tuning curves for the full and sustained phases are very similar – the negative transient response component adds only a weak negative bias to the tuning curves.
**D** Full vs. sustained responses for all stimuli tested in males (gray) and females (magenta). Both measures yield highly correlated responses (Spearman rank correlation r=1.0, p=0). The purely negative transient response component in the full responses adds a negligible negative bias of -0.05mm/s.
All behavioral data from wild type flies of the NM91 strain.

#### Figure S2

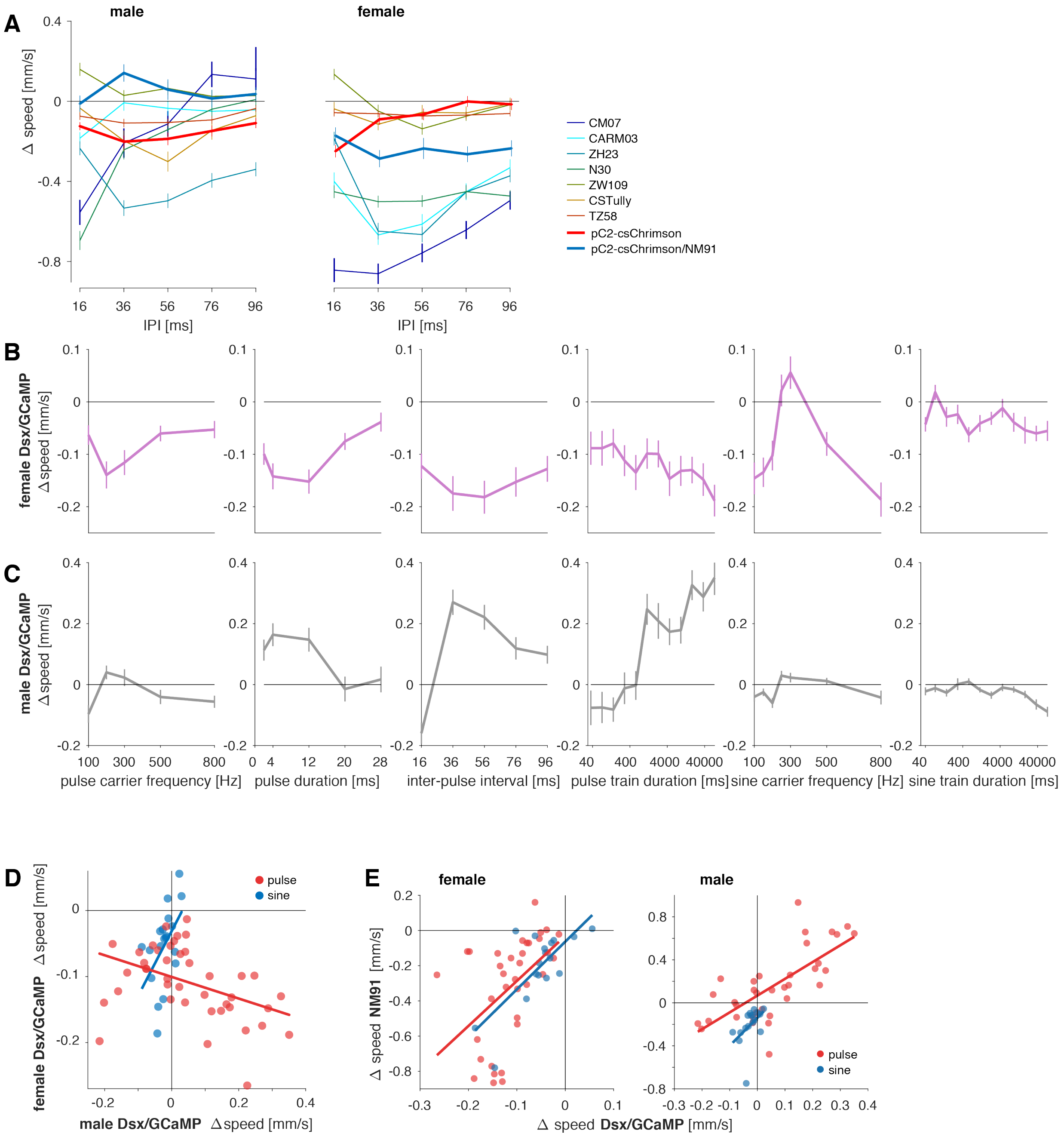

A
male
female
0.4
0
-0.4
-0.8
Δ speed [mm/s]
16 36 56 76 96
IPI [ms]
CM07
CARM03
ZH23
N30
ZW109
CSTully
TZ58
pC2-csChrimson
pC2-csChrimson/NM91
B
female Dsx/GCaMP Δ speed [mm/s]
0.1
0
-0.1
-0.2
C
male Dsx/GCaMP Δ speed [mm/s]
0.4
0.2
0
-0.2
100 300 500 800
pulse carrier frequency [Hz]
0 4 12 20 28
pulse duration [ms]
16 36 56 76 96
inter-pulse interval [ms]
40 400 4000 40000
pulse train duration [ms]
100 300 500 800
sine carrier frequency [Hz]
40 400 4000 40000
sine train duration [ms]
D
female Dsx/GCaMP Δ speed [mm/s]
male Dsx/GCaMP Δ speed [mm/s]
pulse
sine
E
female
male
Δ speed NM91 [mm/s]
Δ speed Dsx/GCaMP [mm/s]

**Figure S2. Related to Figure 2.**

**A** IPI tuning in seven additional strains is diverse and not consistent with the species-typical tuning (male left, female right, see legend for strains used). Note, however, that most strains still respond sex-specifically to pulse song sex, similar to NM91 and Dsx/GCaMP. These strains produce similar responses to song in a natural courtship assay, suggesting that these strains require sensory cues that are missing in FLyTRAP for expressing their song. pC2-csChrimson and pC2-csChrimson/NM91 are the two strains used for optogenetic activation of pC2 in FLyTRAP (see Fig. 5G-L).

**B, C** Locomotor tuning curves for females (C, magenta) and males (D, grey) of the Dsx/GCaMP strain for 6 different features of pulse and sine song. Lines and error bars correspond to the mean±s.e.m. across flies (see Table S1 for a description of all stimuli and N flies). Tuning is similar to that of the wild types strain NM91 (compare Figs. 2A, B).

**D** Changes in speed for Dsx/GCaMP males and females for all pulse (red) and sine (blue) stimuli tested (data from B, C and not shown, see Table S1). Responses to sine stimuli are positively correlated between sexes (r=0.48, p=$3\times10^{-3}$). Pulse responses are negatively correlated (r=-0.58, p=0.01) (compare Fig. 2D).

**E** Comparison of male (left) and female (right) tuning of the NM91 vs. Dsx/GCaMP. Responses for both sine (red) and pulse (blue) are similar between both strains in both sexes (females: pulse r=0.50, p=0.04, sine r=0.59, p=$3\times10^{-4}$; males: pulse r=0.68, p=$4\times10^{-3}$, sine r=0.75, p=$9\times10^{-7}$).

Graphs in A-C show mean ± s.e.m. over individuals (90-150 flies per strain and sex). All correlation values are Spearman's rank correlation. Blue and red lines in D and E correspond to linear fits to the responses to sine and pulse song, respectively. All correlation values are Spearman's rank correlation.

### Figure S3

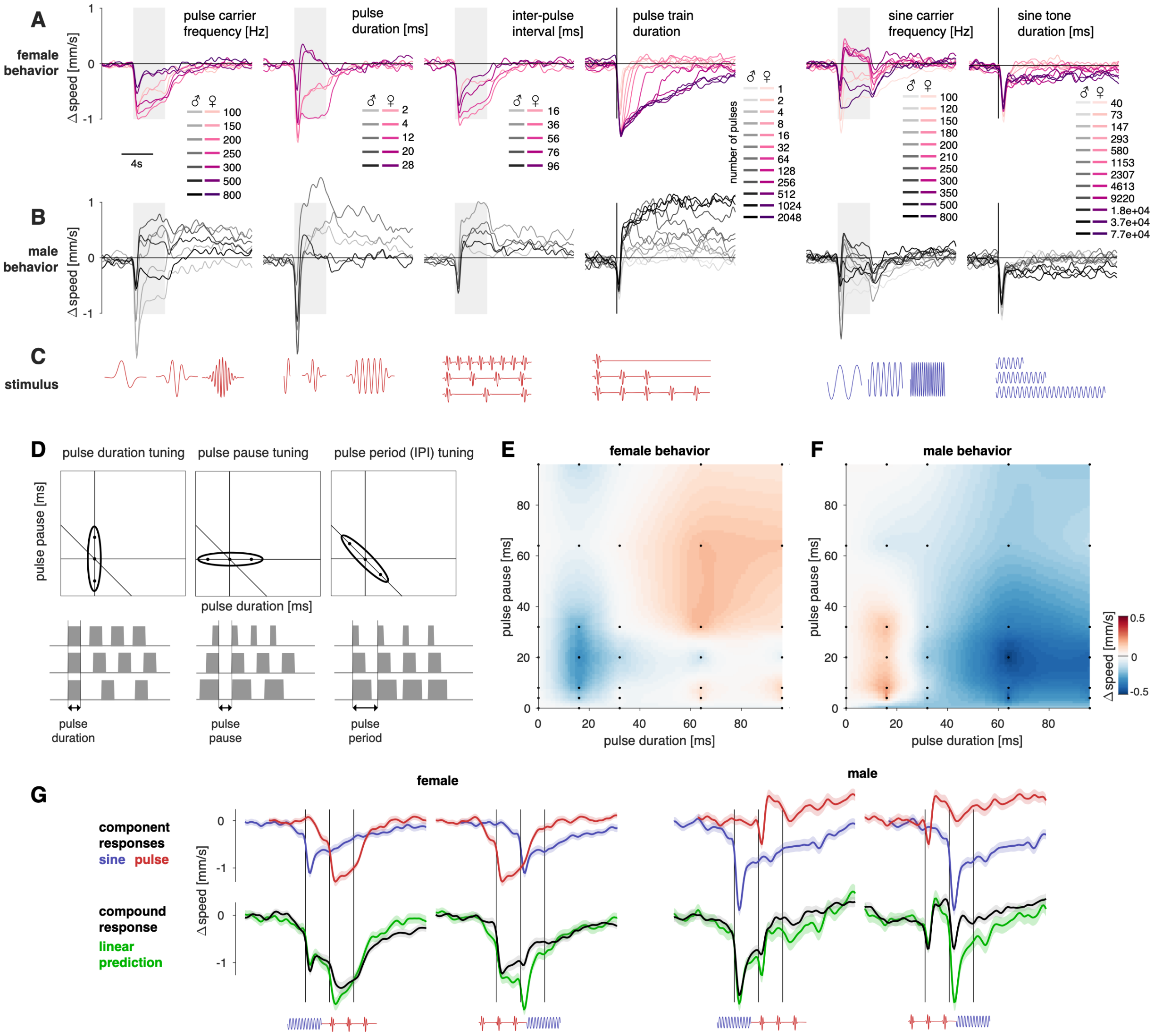

A
female behavior
pulse carrier frequency [Hz]
pulse duration [ms]
inter-pulse interval [ms]
pulse train duration
sine carrier frequency [Hz]
sine tone duration [ms]
Δ speed [mm/s]
4s
number of pulses
B
male behavior
C
stimulus
D
pulse duration tuning
pulse pause tuning
pulse period (IPI) tuning
pulse pause [ms]
pulse duration [ms]
pulse duration
pulse pause
pulse period
E
female behavior
F
male behavior
pulse duration [ms]
pulse pause [ms]
Δ speed [mm/s]
G
female
male
component responses
sine pulse
compound response
linear prediction

**Figure S3. Related to Figure 3.**

**A, B** Locomotor response traces for all stimuli in Fig. 2A, B for females (A) and males (B). Stimulus values are color coded (see legends). Gray shaded areas mark the duration of the sound stimulus. Vertical black lines indicate sound onset for stimulus sets with varying durations (pulse train duration, sine tone duration). Lines correspond to the mean over typically ~120 animals (see supp. table 1 for exact values). Error bars are similar to those in 1C and omitted for clarity.

**C** Pictograms (not to scale) illustrating each song feature examined in A-B. Pulse and sine song features are marked red and blue, respectively.

**D** Three principal types of tuning are observable when testing behavioral responses for stimuli with different pulse durations and pauses. Black ellipses indicate the range of stimulus parameters that evoke strong behavioral responses, dots correspond to the three (gray) stimuli shown below each tuning field. Horizontal, vertical, and anti-diagonal lines mark stimuli with constant pause, duration and period, respectively. Pulse duration tuning (left) corresponds to high selectivity (narrow tuning) for pulse duration and higher tolerance for pulse pauses. Pause duration tuning (middle), corresponds to high selectivity for pulse pause and high tolerance for pulse pause. Note that for both types, the tuning for pulse duration and pulse pause does not interact – e.g. the preferred pause does not change with pulse duration. Pulse period (a.k.a. inter-pulse interval) tuning (right) corresponds to a preference for stimuli with a constant pulse period, given by the sum of pulse duration and pause. For this type of tuning, pulse duration and pulse pause interact – the preferred pulse pause increases with decreasing pulse duration.

**E, F** Locomotor responses of females (E) and males (F) for pulse trains with different pulse durations and pulse pauses. Speed values are color coded (see color bar in F). Black dots mark the parameter combinations of the stimuli tested in FLyTRAP. Intermediate values were obtained using interpolation (see methods). Male and female response fields are similar except for the sign – were females tend to slow (blue colors), males tend to accelerate (red colors), and vice versa (compare Fig. 2C). Responses are more selective for pulse duration than for pulse pause and pulse duration tuning is relatively independent of the pulse duration.

**G** Responses to sequences of sine tones (blue) and pulse trains (red). First and third row correspond to 2s sine followed by 2s pulse for females and males, respectively. For the second and fourth the order is reversed – stimuli start with 2s pulse song and transition into 2s sine song. The top row shows responses to the individual components of the sequence aligned to the onset of the component in the combined stimulus. The bottom row shows responses for both sexes to the compound stimuli (black) and the linear prediction obtained by summing the responses to the individual components (green). The linear prediction matches the measured responses well except at the transition due to transient response to sound onset only present in individual component responses. Lines and shaded areas correspond to the mean ± s.e.m. over 189 female and 217 male flies.

All behavioral data from wild type flies of the NM91 strain.

### Figure S4

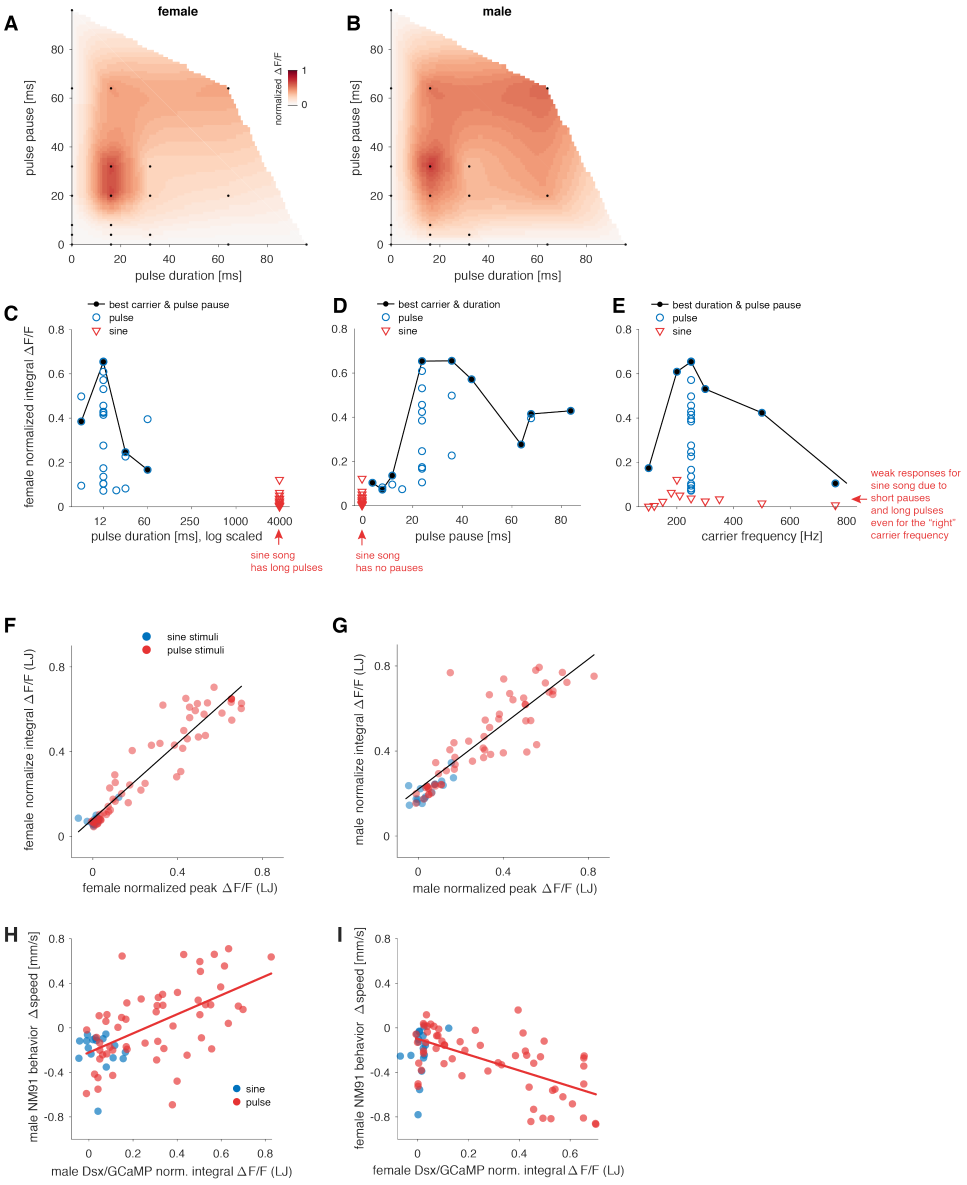

A
female
B
male
normalized ΔF/F
1
0
pulse pause [ms]
pulse duration [ms]
C
best carrier & pulse pause
pulse
sine
female normalized integral ΔF/F
pulse duration [ms], log scaled
sine song
has long pulses
D
best carrier & duration
pulse
sine
pulse pause [ms]
sine song
has no pauses
E
best duration & pulse pause
pulse
sine
carrier frequency [Hz]
weak responses for
sine song due to
short pauses
and long pulses
even for the "right"
carrier frequency
F
sine stimuli
pulse stimuli
female normalize integral ΔF/F (LJ)
female normalized peak ΔF/F (LJ)
G
male normalize integral ΔF/F (LJ)
male normalized peak ΔF/F (LJ)
H
male NM91 behavior Δspeed [mm/s]
male Dsx/GCaMP norm. integral ΔF/F (LJ)
sine
pulse
I
female NM91 behavior Δspeed [mm/s]
female Dsx/GCaMP norm. integral ΔF/F (LJ)

**Figure S4. Related to Figure 3.**

**A, B** Calcium responses from the female (D) or male (E) LJ for pulse trains with different combinations of pulse pauses and pulse durations. The stimuli constitute a subset of those measured in the behavior. LJ tuning for pulse trains with different pulse pauses and pulse durations recapitulates the behavioral tuning (compare Fig. S3E, F): LJ responses are more selective for pulse duration and the preferred pulse duration does not change with pause duration.

**C** Calcium response of the female LJ as a function of the pulse duration for all stimuli in our data set with a duration of 4 seconds. Blue circles correspond to pulse stimuli and red triangles mark sine stimuli, which by definition do not have pauses and can be thought of as very long pulses. Note that besides pulse duration, all stimuli also differ in pulse pause duration and carrier frequency – for instance all sine stimuli (red triangles) differ in carrier frequency. Stimuli with the same pulse duration evoke diverse calcium responses (integral ΔF/F) since they differ in these other stimulus features. The black line connects stimuli that have the optimal pulse carrier frequency (250 Hz) and pulse pause (20 ms). To account for differences in ΔF/F across individuals, values were normalized by the maximal ΔF/F for each individual.

**D** Same calcium response data as in A but now plotted as a function of pulse pause. Sine stimuli correspond to pulse trains with no pauses – they are by definition continuous oscillations. The black line connects stimuli with the optimal pulse duration (12ms) and pulse carrier frequency (250 Hz).

**E** Same calcium response data as in A but now plotted as a function of pulse carrier frequency. Sine stimuli differed in their carrier frequencies. The black line connects stimuli with the optimal pulse duration (12ms) and pulse pause duration (24 ms).

**F, G** Comparison of peak and integral ΔF/F values from the LJ for all stimuli tested in females (F) and males (G). Pulse and sine song are marked with red and blue, respectively. The black lines correspond to the best linear fit. Both measures of calcium responses are highly correlated (males: r=0.93, $p=1x10^{-31}$, female: r=0.94, $p=7x10^{-35}$).

**H, I** Comparison of behavioral and neuronal tuning (LJ) in males (H) and females (I). Neuronal responses are from Dsx/GCaMP flies and behavioral data come from wild type flies (NM91). Dots correspond to the average normalized integral ΔF/F and Δspeed over individuals, lines indicate linear fits. The pattern of correlations is similar to using behavioral data from Dsx/GCaMP flies (compare Fig. 3H,I; males: pulse (red) r=0.58, $p=7x10^{-3}$; sine (blue) r=-0.90, $p=5x10^{-3}$; females: pulse (red) r=-0.66, $p=6x10^{-7}$; sine (blue) r=0.11, p=0.55).

All ΔF/F values from flies expressing GCaMP6m in all Dsx+ cells. All correlation values are Spearman's rank correlation.

### Figure S5

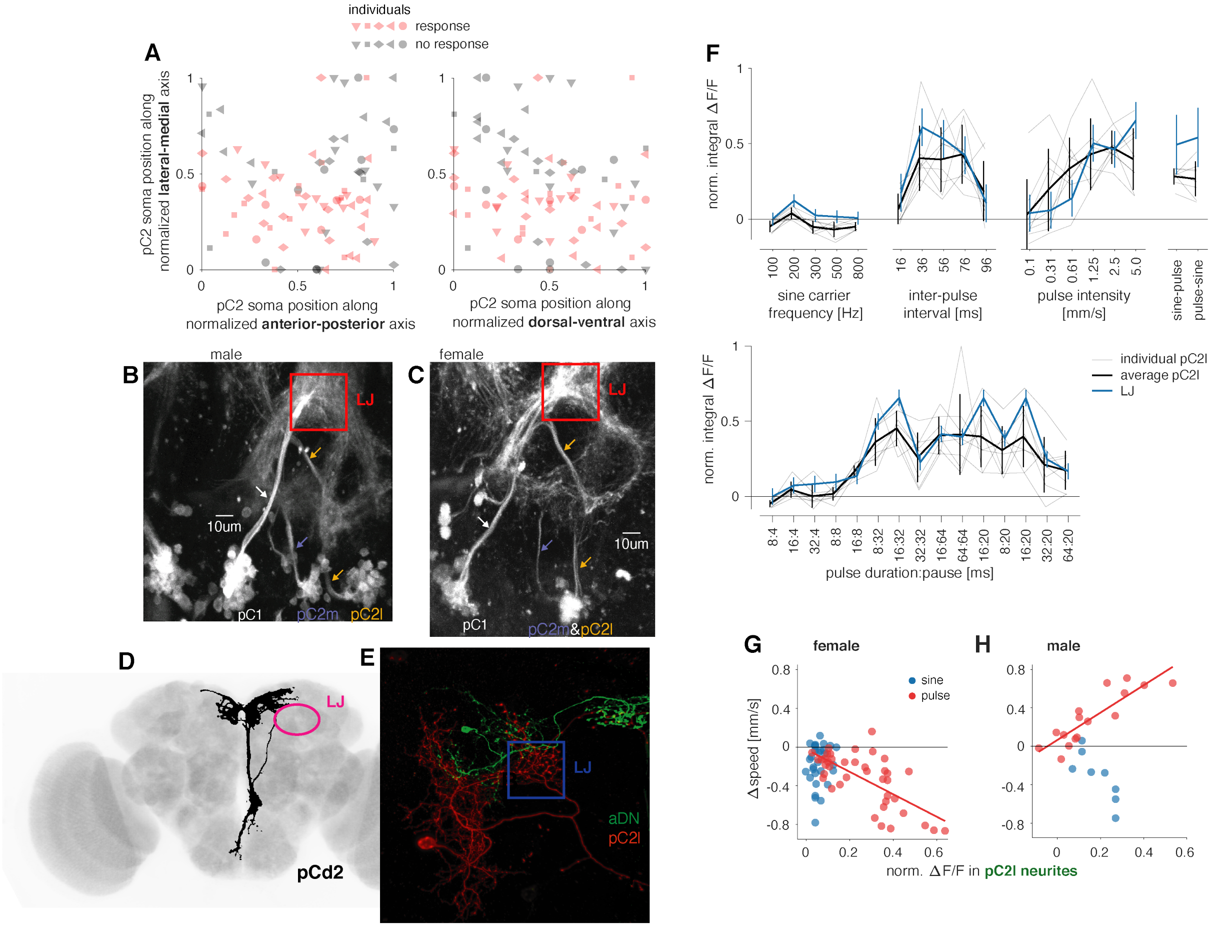

A
individuals
response
no response
pC2 soma position along normalized lateral-medial axis
pC2 soma position along normalized anterior-posterior axis
pC2 soma position along normalized dorsal-ventral axis
B
male
LJ
10um
pC1
pC2m
pC2l
C
female
LJ
10um
pC1
pC2m&pC2l
D
LJ
pCd2
E
LJ
aDN
pC2l
F
norm. integral ΔF/F
sine carrier frequency [Hz]
inter-pulse interval [ms]
pulse intensity [mm/s]
sine-pulse
pulse-sine
individual pC2l
average pC2l
LJ
pulse duration:pause [ms]
G
female
sine
pulse
Δspeed [mm/s]
H
male
norm. ΔF/F in pC2l neurites

**Figure S5. Related to Figure 4.**

**A** Locations of pC2 somas in the female brain with (pink) or without (grey) auditory responses along the lateral-medial and anterior-posterior (left) or dorsal-ventral (right) axis. Soma positions were normalized to between 0 and 1 for each animal. Individual symbols correspond to somas from different animals. Responsive pC2 somas are concentrated in the lateral proportion of the cell cluster (lower half of each plot) and largely correspond to the pC2l subcluster, although we occasionally also observed auditory responses in pC2m somas (see Movie S5).

**B, C** Max z-projection of baseline fluorescence values from a two-photon volumetric scan of a male (B) and female (C) brain expressing GCaMP6m in all Dsx+ neurons. Each of the three clusters (pC1 - white, pC2m – blue, pC2l - yellow) connects to the LJ (red) via unique neurites (arrows). pC2l and pC2m are hard to distinguish by soma location in females since they are intermingled.

**D** A single pCd2 neuron labeled using MCFO (black) registered to a template brain (JFRC2, gray). The lateral junction (LJ) is marked in magenta. pCd1 (not shown, Kimura et al. (2015) and pCd2 do not project to the LJ.

**E** Max z-projection of a confocal stack in which aDN (green) and pC2l (red) were labelled with different colors using MCFO. pC2l but not aDN projects to the LJ (blue).

**F** Tuning of individual pC2 somas (grey), the pC2 population (black), and the LJ (blue) for different song features. Lines and error bars indicate mean±std over 8 different somata recorded from 8 flies.

**G, H** Comparison of calcium responses in the pC2l neurites of Dsx/GCaMP flies and the speed of NM91 flies (males – G, females – H). Correlations are similar when comparing behavioral and neuronal tuning within the same genotype (Dsx/GCaMP, see Fig. 4I, J) (female (G): pulse: r=-0.68, $p=1x10^{-6}$, sine: r=0.15, p=0.44; male (H): pulse: r=0.87, $p=1.5x10^{-5}$, sine: r=-0.82, p=0.01). Each point corresponds to an individual stimulus (Δspeed: N~120 flies per stimulus, ΔF/F: N=10-24 female and 1-6 male flies/stimulus).

All ΔF/F values from flies expressing GCaMP6m in all Dsx+ cells. All correlation values are Spearman's rank correlation.

### Figure S6

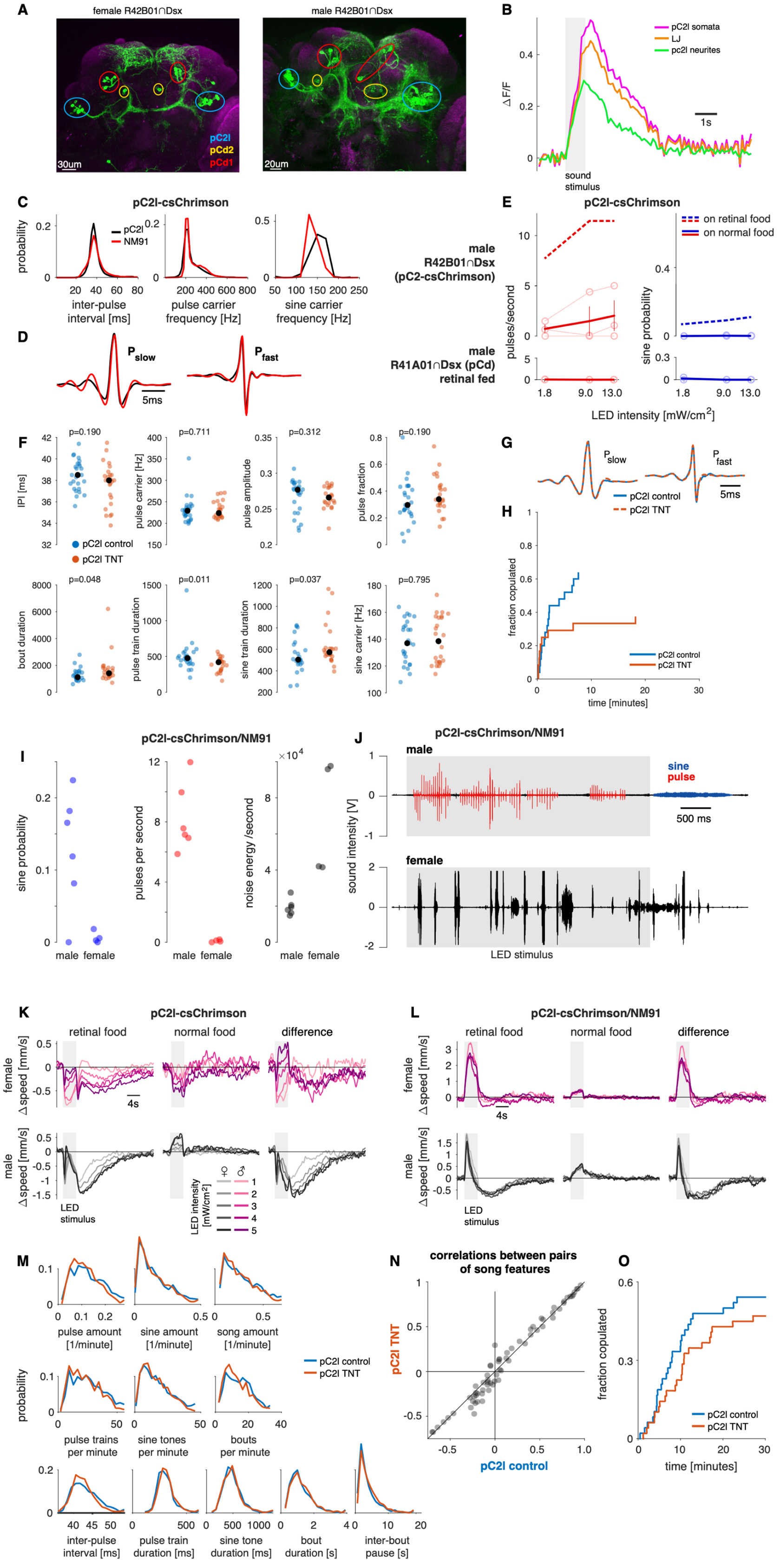
A
female R42B01∩Dsx
male R42B01∩Dsx
pC2l
pCd2
pCd1
30um
20um
B
pC2l somata
LJ
pc2l neurites
ΔF/F
1s
sound stimulus
C
pC2l-csChrimson
pC2l
NM91
probability
inter-pulse interval [ms]
pulse carrier frequency [Hz]
sine carrier frequency [Hz]
D
P slow
P fast
5ms
male R42B01∩Dsx (pC2-csChrimson)
male R41A01∩Dsx (pCd) retinal fed
E
pC2l-csChrimson
on retinal food
on normal food
pulses/second
sine probability
LED intensity [mW/cm2]
F
p=0.190
p=0.711
p=0.312
p=0.190
IPI [ms]
pulse carrier [Hz]
pulse amplitude
pulse fraction
pC2l control
pC2l TNT
p=0.048
p=0.011
p=0.037
p=0.795
bout duration
pulse train duration
sine train duration
sine carrier [Hz]
G
P slow
P fast
5ms
pC2l control
pC2l TNT
H
fraction copulated
pC2l control
pC2l TNT
time [minutes]
I
pC2l-csChrimson/NM91
sine probability
pulses per second
noise energy /second
male
female
J
pC2l-csChrimson/NM91
male
sine
pulse
500 ms
sound intensity [V]
female
LED stimulus
K
pC2l-csChrimson
retinal food
normal food
difference
female Δspeed [mm/s]
4s
male Δspeed [mm/s]
LED stimulus
LED intensity [mW/cm2]
L
pC2l-csChrimson/NM91
retinal food
normal food
difference
female Δspeed [mm/s]
4s
male Δspeed [mm/s]
LED stimulus
M
pulse amount [1/minute]
sine amount [1/minute]
song amount [1/minute]
pC2l control
pC2l TNT
probability
pulse trains per minute
sine tones per minute
bouts per minute
inter-pulse interval [ms]
pulse train duration [ms]
sine tone duration [ms]
bout duration [s]
inter-bout pause [s]
N
correlations between pairs of song features
pC2l TNT
pC2l control
O
fraction copulated
pC2l control
pC2l TNT
time [minutes]

**Figure S6. Related to Figure 5.**
**A** Expression pattern of CsChrimson.mVenus in the intersection of R42B01 and Dsx (green) for females (left) and males (right). Neuropil is labelled with nc82 (magenta). The intersection labels 11/22 female and 22/36 male pC2l neurons, as well as 5-6 pCd1 and 2 pCd2 in either sex
**B** Calcium responses from a female of this line for pulse trains (IPI 36ms) in pC2l somas (blue), the pC2l neurites (yellow) and the LJ (red). We detected auditory response in all 5 pC2l cells visible in the imaging plane, as well as in the LJ and in the pC2l neurites.
**C, D** Song evoked by 655nm activation of R42B01∩Dsx neurons (black, N=7 flies) resembles the natural song produced by wild type flies (NM91, N=47 flies) during courtship (red). Shown are distributions of IPIs, pulse carrier frequencies, and sine carrier frequencies (C) as well as average pulse shapes (D).
**E** Pulse rates and sine song probability upon optogenetic activation of males expressing CsChrimson (red-shifted channel rhodopsin) in R42B01∩Dsx (see A) on food without added retinal (top, solid lines, N=3 flies) or on food with retinal (top, dashed lines, N=7 flies),and males expressing CsChrimson in R41A01∩Dsx (labels pCd1) fed retinal food (middle, N=2 flies). Optogenetic activation of pCd1 in the male (middle) does not evoke song, demonstrating that the singing evoked by R42B01∩Dsx activation in males is due to pC2 and not the pCd1 also labeled in this line. Males expressing CsChrimson in R42B01∩Dsx that were kept on normal food produced some song upon activation, though much less than males fed retinal (see dashed lines). This residual activation likely stems from small amounts of retinal in normal food.
**F** Song features for males in which synaptic output of pC2 (and pCd) was suppressed by expressing TNT in R42B01∩Dsx (pC2/csChrimson) vs. control males. Males courted wild type NM91 females. P-values come from a two-sided rank sum test. There are no statistically significant differences between experimental and control males in any of the shown song features after correcting for multiple comparisons using the Bonferroni method. The features that differ significantly are shown in Fig. 5F. N=25 and 24 males for pC2l control and pC2l TNT.
**G** Average waveforms of the two pulse types produced by *Drosophila melanogaster* males. Same genotypes as in F. Suppressing the synaptic output of pC2l does not affect pulse shapes.
**H** Copulation rates for the males in F. There is a weak but statistically not significant effect of male genotype on copulation rates (p=0.12, Cox's proportional hazards regression model).
**K, L** Speed traces for female (magenta, top) and male (grey, bottom) flies expressing CsChrimson in R42B01∩Dsx using two different genotypes: pC2/csChrimson (K) and pC2/csChrimson/NM91 (L) (see Methods). Shown are responses for flies fed retinal (left), normal food (middle) and the difference between the traces of retinal and normally fed flies (right). Colors correspond to different driving voltages of an array of 655nm LEDs (see legend in K). Lines in K correspond to averages over 70 females and 83 males fed retinal food, and 29 female and 34 males fed normal food. Lines in L correspond to averages over 67 females and 53 males fed retinal food, and 68 female and 84 males fed normal food.
**I** Amount of sine song (left), pulse song (middle), and overall noise (right) produced by males and females upon activation of R42B01∩Dsx in the pC2/csChrimson/NM91 background. Noise energy was calculated as the integral root-mean-square voltage. All values correspond to the signals produced in the 6 seconds following the onset of the 4 seconds of activation.

**J** Examples traces of the signals evoked by activating R42B01∩Dsx in the pC2/csChrimson/NM91 background. Males (top) mainly produce pulse song (red) during the activation and sine song (blue) after activation. Females (bottom) mainly produce noisy signals that do not resemble any of the know modes of courtship song.
**M** Statistics of male song do not change when they court pC2l TNT females (using the R42B01∩Dsx driver). Shown are distributions of 11 song parameters from NM91 males courting pC2 control (blue) or pC2 TNT females (orange).
**N** Female genotype does not change correlation statistics (correlations between pairs of song parameters) in the song of NM91 males. Shown are all unique pair-wise correlations between the 11 song parameters in F for the song of NM91 males courting pC2l control (x values) or pC2l TNT females (y values).
**O** Cumulative fraction of copulated pairs of NM91 males courting pC2l control (blue) or pC2l TNT females (orange). There is a weak but statistically not significant effect of female genotype on copulation rates (p=0.19, Cox's proportional hazards regression model).
M-O: Data from 48 pC2 control and 48 pC2 TNT pairs.

**Figure S7**

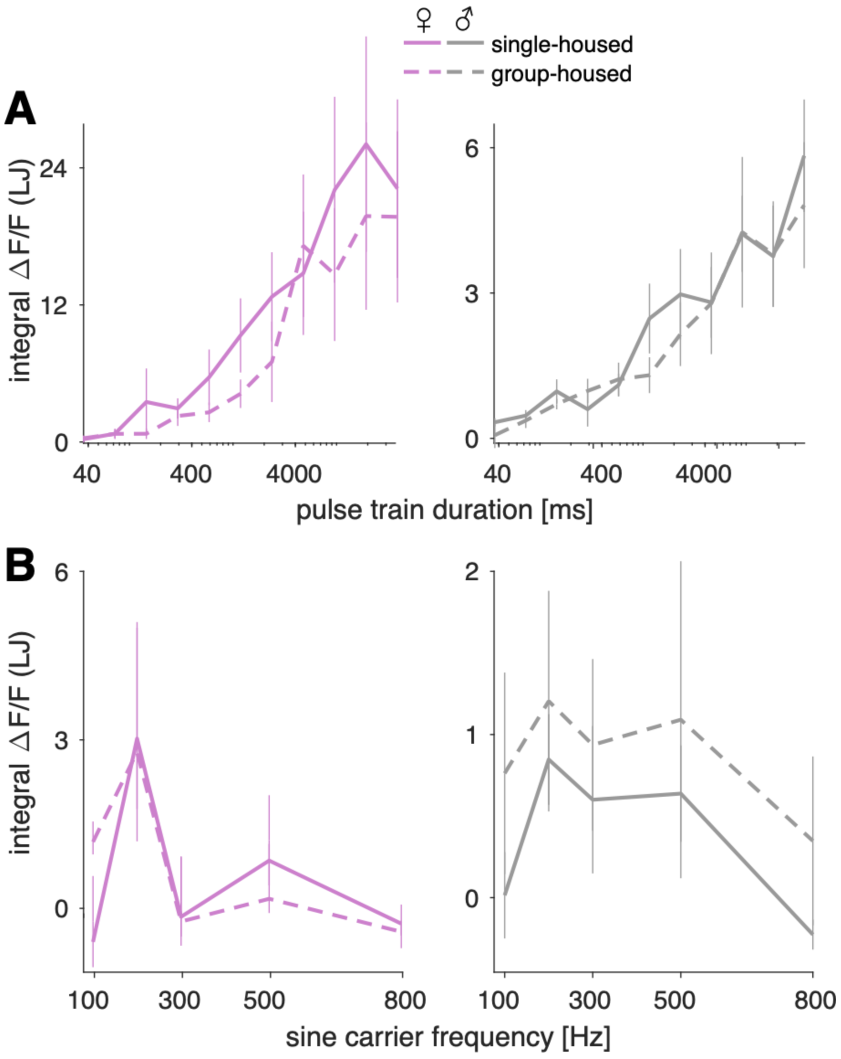

♀ ♂
single-housed
group-housed
A
integral ΔF/F (LJ)
24
12
0
40
400
4000
6
3
0
40
400
4000
pulse train duration [ms]
B
6
3
0
integral ΔF/F (LJ)
100
300
500
800
2
1
0
100
300
500
800
sine carrier frequency [Hz]

**Figure S7. Related to Figure 6.**

**A** Calcium responses in pC2 neurons (measured via the LJ) for pulse trains with different durations (IPI=36ms) in single- and group-housed (solid and dashed lines) females (left) and males (right). Data from 5-6 flies for each condition (single/group-housed, females/males).

**B** Same as A, but for sine carrier frequency. Data from 20/5 single/group-housed females and 12/6 single/group-housed males. Responses to pulse trains with different durations and to sine tones with different carrier frequencies do not change substantially with housing conditions. Lines and error bars correspond to mean ± s.e.m over flies.

All $\Delta F/F$ values from flies expressing GCaMP6m in all Dsx+ cells.

**Movie legends**

**Movie S1** – Female flies (wild type NM91) in FLyTRAP. Track histories are shown as colored trails. Movie speed corresponds to real time.

**Movie S2** – Males extend their wings in response to pulse song in FLyTRAP. In this example, 10 out of 12 flies extended their wings after stimulus onset (4 second pulse train, inter-pulse interval = 36ms). Responding flies are marked with red circles. Movie is slowed down 4X. Some flies extend their wings spontaneously (not triggered by sound), see for example fly 5 in this movie.

**Movie S3** – Two-photon Calcium imaging of the female lateral junction and pC2l neurites. GCaMP6m is expressed in all the Dsx+ neurons. The response in the lateral junction (LJ) and pC2l neurites are highly correlated (see also Fig 4F). Three responses are shown for a single fly – 4 second pulse trains with inter-pulse intervals of 16/36/76ms. The text in the corner appears when the sound stimulus is on. Movie speed corresponds to real time.

**Movie S4** - Two-photon Calcium imaging of the male lateral junction and pC2l neurites (same as in S3).

**Movie S5** – Two-photon Calcium imaging (single plane) of female pC2 cell bodies. The responses to a 4 second pulse train and to a 4 second sine tone (carrier frequency – 200Hz) are shown. In most females the pC2l and pC2m cell bodies can not easily be separated into distinct clusters. Movie speed corresponds to real time.

**Movie S6** – Two-photon Calcium imaging (single plane) of male pC2 cell bodies (same stimuli as in movie S5). pC2l and pC2m cell bodies can be spatially organized in two distinct clusters. Movie speed corresponds to real time.

**Movie S7** – Optogenetic activation of pC2l neurons in an isolated male. Three light intensities are shown (1.8, 9, 13 mW/cm2). Sound is recorded using microphones that tile the chamber floor. Video and audio are synced and played at real time.
